# Navigating “tip fog”: Embracing uncertainty in tip measurements

**DOI:** 10.1101/2024.08.19.608647

**Authors:** Jeremy M. Beaulieu, Brian C. O’Meara

**Author notes:** **Data availability.** All scripts and substantial outputs are available at https://github.com/thej022214/MeasurementError and https://zenodo.org/doi/10.5281/zenodo.13685561. **Author contributions:** J.M.B. and B.C.O. contributed equally. **Conflict of Interest Statement:** The authors declare that they have no conflict of interests.

## Abstract

Nature is full of messy variation, which serves as the raw material for evolution. However, in comparative biology this variation is smoothed into averages. Overlooking this variation not only weakens our analyses but also risks selecting inaccurate models, generating false precision in parameter estimates, and creating artificial patterns. Furthermore, the complexity of uncertainty extends beyond traditional “measurement error,” encompassing various sources of intraspecific variance. To address this, we propose the term “tip fog” to describe the variance between the true species mean and what is recorded, without implying a specific mechanism. We show why accounting for tip fog remains critical by showing its impact on continuous comparative models and discrete comparative and diversification models. We rederive methods to estimate this variance and use simulations to assess its feasibility and importance in a comparative context. Our findings reveal that ignoring tip fog substantially affects the accuracy of rate estimates, with higher tip fog levels showing greater biases from the true rates, as well as affecting which models are chosen. The findings underscore the importance of model selection and the potential consequences of neglecting tip fog, providing insights for improving the accuracy of comparative methods in evolutionary biology.

## Introduction

*True, one portrait may hit the mark much nearer than another, but none can hit it with any very considerable degree of exactness. So there is no earthly way of finding out precisely what the whale really looks like. And the only mode in which you can derive even a tolerable idea of his living contour, is by going a whaling yourself; but by so doing, you run no small risk of being eternally stove and sunk by him. —Herman Melville, Moby Dick*

Messy variation is ubiquitous in nature. This variation provides the raw material for evolution, while smoothed averages of it are crucial for phylogenetic comparative methods. Typically, multiple individuals — or even one — are used to calculate a mean, which is then used as the representative value for a species in trait evolution, correlation, or diversification studies. While we acknowledge some scatter around this idealized value, we often dismiss it as minor, believing it only slightly reduces the power of our methods, if at all. However, ignoring this variation introduces more than just noise; it can fundamentally undermine analyses. It can change the models we choose as best fits to the data (e.g., Sylvestro et al., 2015), produce misleadingly precise parameter estimates (e.g., Ives et al., 2007), and reveal seemingly compelling patterns that are entirely artificial (e.g., O’Meara & Beaulieu, 2024). Ignoring variance at the tips is not just muddying the waters, it is creating spurious new islands towards which we set sail.

The sources of this variation are numerous and additive. First, every measurement carries uncertainty, a concept ingrained in us from the early days of learning science with quizzes about significant digits and finite resolution of tick marks on our meter sticks. Then there is biological uncertainty, such as the exact length of a squid’s tentacle or the precise point where a leaf begins. Sampling uncertainty arises when a subset is used to represent the entire group. Finally, true intraspecific variability exists due to differing optimal conditions, genetic drift within populations, and phenotypic plasticity. Taken together, even the most precise measurement from one population might not reflect the overall species mean.

In comparative biology, we often overlook this complexity of uncertainty. We might consider “measurement error” as the standard error of the observed measurements for species, but the common default is to assume this error is zero. Moreover, the array of factors contributing to this uncertainty extends far beyond what we traditionally categorize as measurement error. This intraspecific variance has been described with various terms, including “specific variances” (Cheverud et al., 1985), “residual variation” (Lynch, 1991), “phenotypic variation” (Felsenstein, 2008), “measurement variance” (Labra et al., 2009; Hansen & Bartroszek), and “measurement error” (Harmon & Losos, 2005; O’Meara et al., 2006; Silvestro et al., 2015). Some of these terms suggest specific mechanisms. Other terms are more descriptive but may have meanings outside our field that can cause confusion. To avoid ambiguity, we propose a new term: “tip fog.” This term captures the variance that occurs at the present between the true species mean derived from the evolutionary process and what an experimenter records as a value, without being tied to any particular mechanism, and it applies to characters that are discrete or continuous.

Here, we show why accounting for tip fog is critical by showing its impact on both continuous and discrete comparative models. We rederive methods to estimate this variance and assess its feasibility and importance in a comparative context. By highlighting the consequences of ignoring tip fog, we urge the community to adopt these methods as default standards into all comparative analyses to avoid the pitfalls of misleading model selection, inferring biased parameter estimates, and interpreting artificial patterns.

## Estimating tip fog for continuous trait evolution

There has been extensive research on the importance of tip fog (under various names) in continuous models, including approaches like independent contrasts (Felsenstein, 2008) and univariate models where model parameters or the comparison of model fit are relevant (Harmon & Losos 2005; Ives et al., 2007; Revell & Reynolds, 2012; Silvestro et al., 2015). All these studies concluded that tip fog can significantly affect the results and recommend its inclusion in analyses, a point particularly emphasized by Silvestro et al. (2015). However, its use remains remarkably uncommon. Popular software packages, such as *ouch* (Butler & King, 2004), *surface* (Ingram & Mahler, 2013), and *revBayes* (Hohna et al., 2016), do not allow accounting for tip fog in their Ornstein-Uhlenbeck models. Packages like *OUwie* (Beaulieu et al., 2012), *bayou* (Uyeda & Harmon, 2014), *SLOUCH* (Labra et al., 2009); and *Rphylopars* (Goolsby et al., 2016) allow specifying tip fog as the standard error of species means but default it to zero; the function fitContinuous in *geiger* (Pennell et al., 2014) also allows it to be inferred, though it also assumes a default value of zero.

The reluctance to incorporate tip fog may stem from the difficulty of obtaining empirical values of sample variance (which is a subset of tip fog) — it is challenging to acquire species means for all traits, let alone the variance in that value. Additionally, any observed value is likely to be an underestimate, capturing only some sources of variation. Estimating tip fog introduces another free parameter into the model, so despite arguments for its potential utility, researchers might opt for simpler models.

The way we parameterize tip fog in *OUwie* is rather straightforward and follows from O’Meara et al. (2006) and Ives et al. (2007). For a standard Brownian motion model, the evolution of a trait over time is modeled as a random walk, with the rate of evolution described by the rate parameter σ^2^. The expected variance-covariance matrix, **V**, reflects the variances and covariances of trait values among different species, based on their shared evolutionary history.

To incorporate tip fog, ζ_*c*_ (“zeta”, to avoid confusion with the other Greek letters populating our models), the within-species variance is added to the diagonal elements of **V**, representing the additional variance in trait values due to factors specific to each species. The modified variance-covariance matrix, **V**\*, is expressed as:

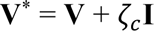

where **I** is the identity matrix. Adding ζ_*c*_ to the diagonal of **V** allows the model to account for both the phylogenetic covariance among species as well as estimate the additional variance within each species. It is possible to have a different ζ_*c*_for each individual species: the variance in log length from measurement ambiguity for a squishy squid is likely larger than the variance in log length for a crunchy crustacean species, for example. One might imagine that a ζ_*c*_ as a percentage of each species’ mean could also work. In our implementation here, we assumed all species had the same value for tip fog when estimating it (but not necessarily when simulating). This approach aligns with the recommendation by Hansen & Bartoszek (2012) for cases where individual species have small sample sizes (see also Labra et al., 2009). Specifically, they assume that within-species variances are similar among species and therefore recommend calculating the within-species variance as a sample size weighted average of each species.

For more complex models, such as those allowing the rate parameter σ^2^ to vary and/or allowing traits to evolve towards specific trait “optima” (i.e., like Ornstein-Uhlenbeck models) based on a discrete regime, the addition of ζ_*c*_ follows the same formulation described above. The only difference lies in how **V** is constructed. However, the impact of not accounting for tip fog on evolution rates in such models is quite dramatic (see results below).

To illustrate, we conducted a set of simulations where the generating models were a multiple-rate Brownian motion model (BMS) and a multiple-optima, multiple-rate Ornstein-Uhlenbeck model (OUMV). We first created an identifiable two-state regime mapping on a randomly generated 200-tip phylogeny (birth rate set to 0.4 events Myr^-1^, and death rate of 0.2 events Myr^-1^) in *TreeSim* (Stadler 2011) with the root to tip length scaled to one. Under both generating models, we set the rate for the first regime to 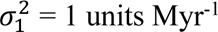 and scaled the rate for the second regime to take on either 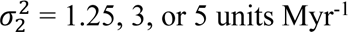. For the OUMV model, we set a global α parameter, which controls the rate of the “pull” towards individual trait means (here we set θ_1_ = 1, and θ_2_ = 3), ensuring that the half-life corresponds to 50% the height of the tree (i.e., α = 1.39 units Myr^-1^).

Tip fog was simulated by resampling each tip value from a normal distribution centered at the individual species mean and with a standard deviation that was a percentage of the mean for each species (which is more complex, but likely more realistic, than assuming the same standard deviation for each species). We generated 100 data sets each, where the percentage varied from 1%, 2.5%, 5%, 10%, to 20% of each individual species mean. Each data set was then evaluated under BM1, BMS, OU1, OUM, and OUMV models assuming three different type of tip fog settings. The first scenario assumed a ζ_*c*_ = 0, meaning species values were treated as known without error, which is the default setting in most programs. In the second scenario, we sampled each species’ mean five times under each tip fog condition and estimated the within-species sample variance as a sample-size-weighted average of the variances for each species (i.e., + User ζ_*c*_). This follows the procedure recommended by Hansen & Bartoszek (2012). However, since the sample sizes are identical, the weighted average variance, 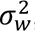, is simply the mean variance across all species, making the sample variance for all species 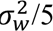. We then converted this value to a standard error by taking the square root and inputting it as a separate data column in the analyses. Finally, we estimated tip fog directly from the species means only as a parameter in the model (i.e., + ζ_*c*_). Estimates for evolutionary rates were summarized as weighted harmonic averages using Akaike weights, which is equivalent to weighted arithmetic means of the wait times; estimates of trait means were based on a weighted arithmetic average. All simulations were performed in the R package *OUwie* (Beaulieu et al., 2012).

The results of these simulations are presented in Figures 1-2, S1-2, and Table S1. Generally, failing to account for tip fog can lead to mistakenly strong support for more complex models. For example, when using the BMS generating model, the OUMV model consistently received over 75% support when tip fog was set at 10% and 20% (Table S1). It can also substantially affect the estimates of evolutionary rates in complex models with multiple rate regimes. As the level of fog increases, the evolutionary rates are increasingly biased upward, regardless of the regime. For example, with greater than 5% fog under a BMS generating model the average rate in regime 1 and 2 is 1.5-to 7-fold higher for all three generating models we tested. In contrast, when ζ_*c*_ is estimated as part of the model (+ζ_*c*_) or provided as an estimate of the within-species variance for each species (+ User ζ_*c*_), the evolutionary rates generally align more closely with their true values (for the same models as above, 0.9 to 1.1 times the true value). However, with a user-supplied estimate of tip fog, when there were large differences in the rate between the two regimes in the OUMV model 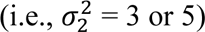, substantial bias occurs at fog levels greater than 10%, affecting the rate estimates in both regimes, ranging from being 1.5 and 3 times the true value (Fig. 1, S1). Estimating tip fog as a parameter within the model is shows very modest biases for both regimes.

**Figure 1.**
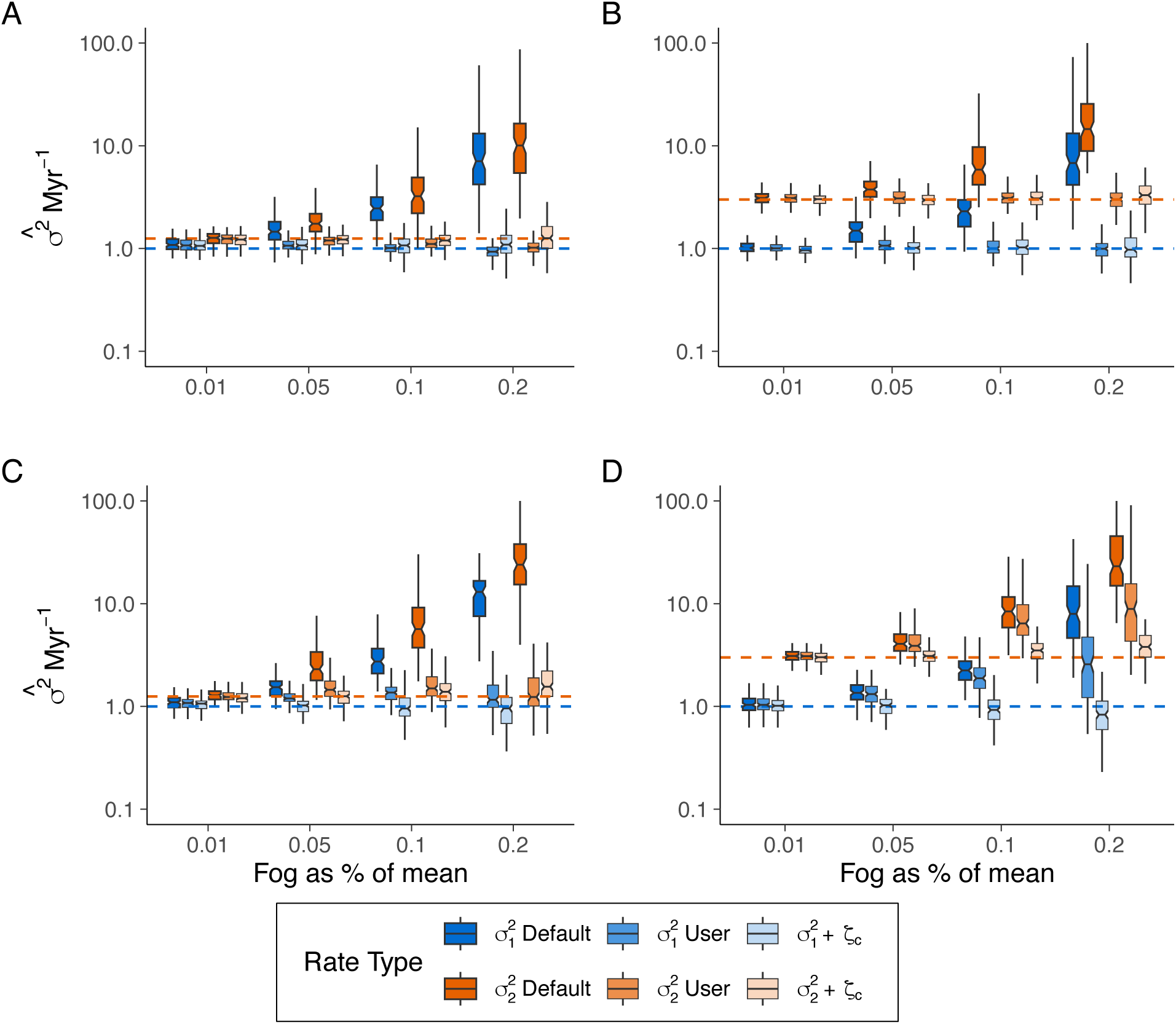
Uncertainty in estimating the evolutionary rate (σ^2^) as a function of tip fog, with the generating model assuming various (**A**, **B**) multiple-rate Brownian motion (BMS) and (**C**, **D**) multiple mean, multiple-rate Ornstein-Uhlenbeck (OUMV) models. Panels (**A**) and (**C**) depict cases where the rate for regime 2 is 1.25-times that of regime 1, while (**B**) and (**D**) show a rate 3-times that of regime 1 (see Fig. S1 for results where the rate for regime 2 is 5-times that of regime 1). Tip fog was simulated by resampling each tip value from a normal distribution centered at the individual species mean and with a standard deviation that was a percentage of the mean. Data sets were then evaluated under BM1, BMS, OU1, OUM, and OUMV models, with rates summarized using a weighted harmonic mean based on Akaike weights (see text). Darker boxes indicate rate summarized across models excluding tip fog (Default), which show an upward bias in evolutionary rates as fog levels increase, regardless of the regime. The less saturated boxes represent rates summarized across models that either calculate tip fog from simulated “measurements” (+User) or estimate tip fog (+ζ_*c*_), where evolutionary rates generally align more closely with true values in nearly all cases (main exception being in (**D**) for +User estimates). Dashed blue and orange lines indicate the generating values for regimes 1 and 2, respectively.

**Figure 2.**
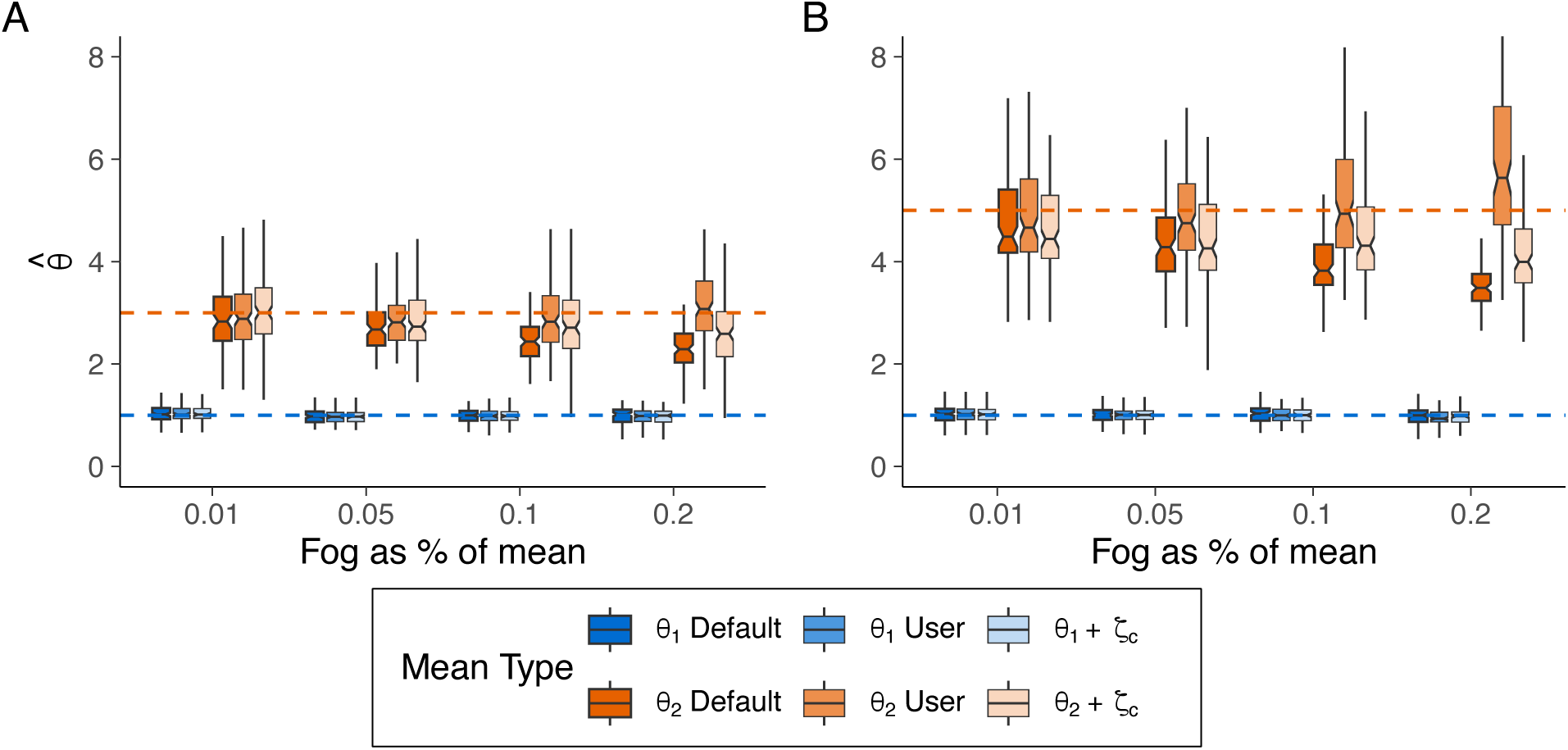
Uncertainty in estimating the trait means (θ*_i_*) when the generating model is a multiple-mean Ornstein-Uhlenbeck model (OUM), and the simulated data sets contained differing levels of tip fog. (**A**) depicts a scenario where the trait mean for regime 2 is 3-times that of regime 1, whereas (**B**) depicts a scenario where regime 2 is 5 times that of regime 1. Tip fog was simulated by resampling each tip value from a normal distribution centered at the individual species mean and with a standard deviation that was a percentage of the mean. Data sets were evaluated under BM1, BMS, OU1, OUM, and OUMV models, with rates summarized using a weighted mean based on Akaike weights (see text). Darker boxes indicate θ*_i_* summarized across models excluding tip fog (Default); the less saturated boxes represent θ*_i_* summarized across models that either calculate tip fog from simulated “measurements” (+ User) or estimated tip fog (+ζ_*c*_) directly from the data. Dashed blue and orange lines indicate the generating values for regimes 1 and 2, respectively. In the Default and +ζ_*c*_ scenarios, as the amount of tip fog increases, estimates for θ_2_ are increasingly underestimated, irrespective of whether tip fog was estimated.

For the OUMV models, tip fog had a pronounced effect on estimates of the trait mean (θ_2_) for regime 2 only, regardless of the underlying rate or whether fog was estimated as part of the model (Fig. S2). Interestingly, incorporating tip fog as the user-supplied estimate of the variance did not generally show this pattern. To better understand this, we conducted an additional simulation, assuming a multiple-optima OU model (OUM) where the evolutionary rate was set equal across regimes σ^2^ = 2 and varied θ_2_ to be 1.25x, 3x, and 5x on a pectinate tree with an identifiable two-state regime mapping. Results from these simulations exhibited a similar pattern: in the presence of fog, the estimates of theta for the second regime are increasingly underestimated as the amount of tip fog increases (Fig. 2). Examining the confidence regions surrounding estimates of θ_2_, α, and σ^2^ using *dentist* (Boyko & O’Meara, 2024) reveals that tip fog introduces greater uncertainty in estimates of α and σ^2^, which incidentally impacts estimates of θ_2_, even when ζ_*c*_ is included in the model (Fig. S3). This uncertainty is also reflected in model support; the presence of fog leads to some support for models that include additional complexity, in particular, support for the OUMV model (Table S1). Overall, these results suggest that any model fit should be interpreted with consideration of the parameter estimates and the underlying uncertainty.

We also analyzed empirical estimates of tip fog from various datasets (see Supplementary Materials). In some cases, particularly for smaller datasets, the best models assumed tip fog was fixed at zero, or in one case, tip fog alone explained the data without the need for an evolutionary model. However, in other datasets, tip fog could be substantial. For example, in Alencar et al. (2024), the log of topographical complexity (across 663 species) had a tip fog estimate of 47% of the trait mean, with tip fog expected to explain 44% of the total trait variance. For log precipitation, the best model had no tip fog, while the next best model (with a ΔAICc of 0.88 which is still substantial weight) estimated tip fog at 8% of the mean, explaining 14% of the total variance (Table S2).

### Impact of tip fog on discrete trait evolutionary models

The impact of character misassignments on the accuracy and reliability of continuous-time Markov models remains largely unexplored outside of tree inference contexts (e.g., Felsenstein, 2004; Ho et al., 2007; Rambaut et al., 2009; Han et al., 2013; Kuhner & McGill, 2014; Davis & Navin, 2016). Misassignments, whether due to data collection errors or legitimate polymorphisms, introduce uncertainty about the true state of a species and can lead to erroneous estimates of transition rates between states and exaggerated biological patterns (Han et al., 2013; O’Meara & Beaulieu, 2024). Here we apply a framework, first proposed by Felsenstein (2004), for continuous-time Markov models that allows for the simultaneous estimation of character misassignments and state transition rates. Conceptually, this approach is best understood as a type of hidden Markov model (HMM). In such models, the true state is not directly observed, making it “hidden,” while the observed data represent a noisy or misclassified version of these states (also see Jackson et al., 2003). The goal of the model is to infer the underlying true states by estimating the likelihood of observing the data given the true states. It is rather straightforward in that the general model remains unchanged except we alter the observed probabilities at the tips.

Suppose that *i* indexes *n* tips in a tree, and that *Si* represents the true underlying state of tip *i* and that *Oi* corresponds to the observed state for tip *i*. When these states are known exactly, as is assumed by any standard continuous-time Markov model, we assume the state of each tip is *P*(*Oi* = *o*| *Si* = *o*) = 1. In the binary case, *o* might represent the presence or absence of a particular character state (e.g., woody versus herbaceous plants, feathers versus no feathers). However, when these observations are subject to uncertainty, as is often the case, we assume then that the observed states, *Oi*, are generated conditionally on the true states, *Si*. The probability of the observed state *o* given that the true state is *s* can be expressed by *P*(*Oi* = *o*| *S_i_* = *s*) = 1 - ζ_o,s_ where ζ_o,s_ defines what we refer to as the “tip fog probability”. Thus, the probability at a given tip then becomes 1 - ζ_o,s_for the observed state and ζ_o,s_ for the alternative state. For an arbitrary number of states when state *i* is observed, the probability of each alternative state (i.e., *j* ≠ *i*) is ζ*_i,j_* with the observed state being 1 − ∑*_j#i_* ζ*_i,j_*, where ζ*_i,j_* represents the probability that the observed state is *i* when the true state is *j*.

The likelihood of the model is obtained by maximizing the standard likelihood formula, *L* = *P*(*D* | **Q**, T, ζ_*d*_), for observing character states, *D*, given the continuous-time Markov model, **Q**, a fixed topology with a set of branch lengths (denoted by *T*). Note that we have added an additional free parameter in the formula, ζ_*d*_, that denotes a single tip fog probability for all observed states. When ζ_*d*_ = 0, the likelihood simply reduces to the likelihood of a standard continuous-time Markov model. We implemented these “tip fog” models as part of the *corHMM* package (Beaulieu et al., 2013; Boyko & Beaulieu, 2021), allowing for the specification of as many tip fog probabilities, ζ*_i_*, as there are unique observed states in the data set.

We conducted a simulation study to assess the impact of ignoring tip fog, as well as the overall behavior when estimating it. A 200-tip phylogeny was generated (birth rate set to 0.4 events Myr^-1^, and death rate of 0.2 events Myr^-1^) in *TreeSim* (Stadler 2011), which was used to simulate 100 datasets assuming equal transitions rates of *q01* = *q10* = 0.025 transitions Myr-1 between binary states. We varied the root age of the phylogeny to take on four ages: 5, 10, 15, and 20 Myr. In a recent publication (O’Meara & Beaulieu, 2024) we showed that inaccuracies in character state assignments significantly affect younger trees more due to shorter overall tree lengths, and so varying clade age was meant to mimic this effect. To simulate tip fog, we randomly altered the observed state of 1%, 5%, 10%, 15%, or 20% of taxa to be the reverse of its true state. For each simulation replicate, we fit two classes of models: 1) equal rates (ER, single rate for all transitions) and all rates different (ARD, two independent rates) without estimating tip fog probabilities (i.e., referred to as “Default”), and 2) ER and ARD with each estimating a single ζ_*d*_ parameter. Rate estimates within each of the model classes were then summarized by calculating a weighted harmonic mean of each transition parameter using the Akaike weights (*wi*).

As with continuous trait data, failing to account for tip fog dramatically inflates transition rates. We found that the magnitude of the upward bias increases as the degree of tip fog also increases (Fig. 3). Even a tip fog probability of 1%, which is expected to result in just two taxa out of 200 being misassigned to the wrong state, substantially biased transition rates upward. As expected, the magnitude of the effect did depend on clade age. For example, with 1% tip fog probability in a 5 Myr tree, rates more than double their true value (*q01* = 0.049 and q*10* = 0.069 transitions Myr^-1^), while for a clade age of 20 Myr, rates are only slightly upward biased (*q01* = 0.030 and q*10* = 0.033 transitions Myr^-1^). Nevertheless, when tip fog was 5% and higher, the model-averaged rates were orders of magnitude higher than the true values regardless of clade age (Fig. 3). When tip fog probability was estimated as part of the model (+ζ_*d*_), the transition rates remain relatively stable across different fog levels (Fig. 3). Although younger clades still show a slight upward bias, it does not systematically increase with increasing tip fog as it does in the default model fits. Notably, estimates of ζ_*d*_ are consistently centered on their true values, indicating the model effectively infers the degree of tip fog present in these data sets (Fig. 3C).

**Figure 3.**
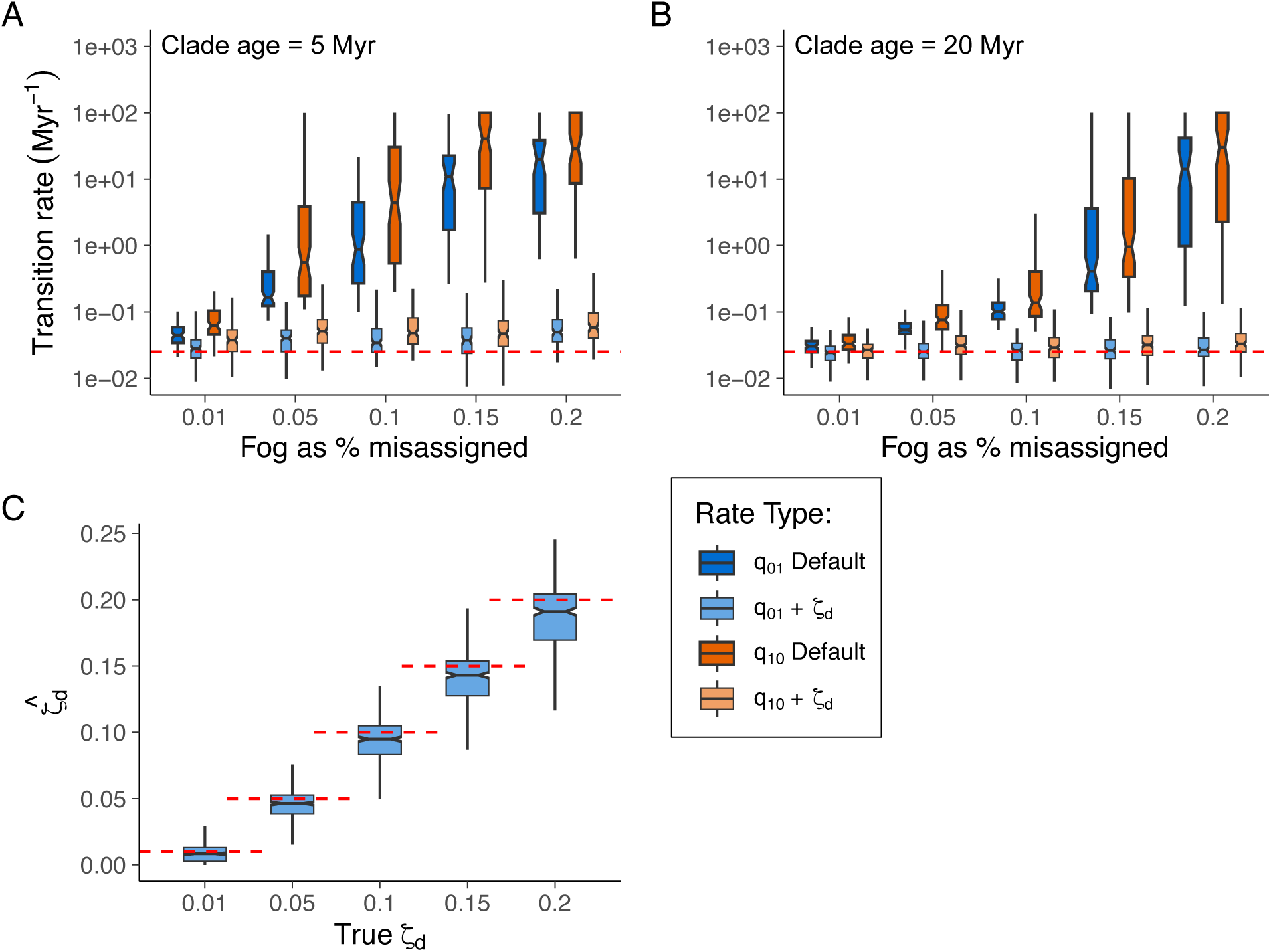
Uncertainty in estimating transition rates using an equal rates (ER) continuous-time Markov model (*q01* = *q10* = 0.025 transitions Myr^-1^) with increasing levels of tip fog. Panels (**A**) and (**B**) show rate uncertainty for clades aged 5 Myr and 10 Myr, respectively. Younger clades show greater uncertainty due to shorter tree lengths (see main text). To simulate tip fog, we randomly altered the observed state of 1%, 5%, 10%, 15%, or 20% of taxa to be the reverse of its true state. Data sets were evaluated under an equal-rates model (ER, single rate for all transitions) and an all-rates different model (ARD, two independent rates). Rate estimates within each of the model classes were then summarized by calculating a weighted harmonic mean of each transition parameter using the Akaike weights. Darker boxes indicate transition rates summarized across models excluding tip fog (Default); the less saturated boxes represent transition rates summarized across models that estimated tip fog (+ζ_*d*_). Dashed blue and orange lines indicate the generating values for state *0* and state *1*, respectively. (**C**) depicts uncertainty in ζ_*d*_ estimates across all clade ages, with the dashed red line indicating the true ζ_*d*_.

We also found that failing to account for tip fog substantially impacts model weights. With just 5% tip fog, the weight shifts almost entirely (>90%) to the ARD model, a pattern that was consistent across all clade ages (Fig. 4; Table S3). When the tip fog probability is estimated under an ER generating model, the Akaike weights favor the generating ER model with the distribution of the weights converging towards the null weight expected based solely on the penalty term (Fig. 4). This indicates that uncorrected tip fog not only inflates rate estimates but can also erroneously make the data choose models that are too complex.

**Figure 4.**
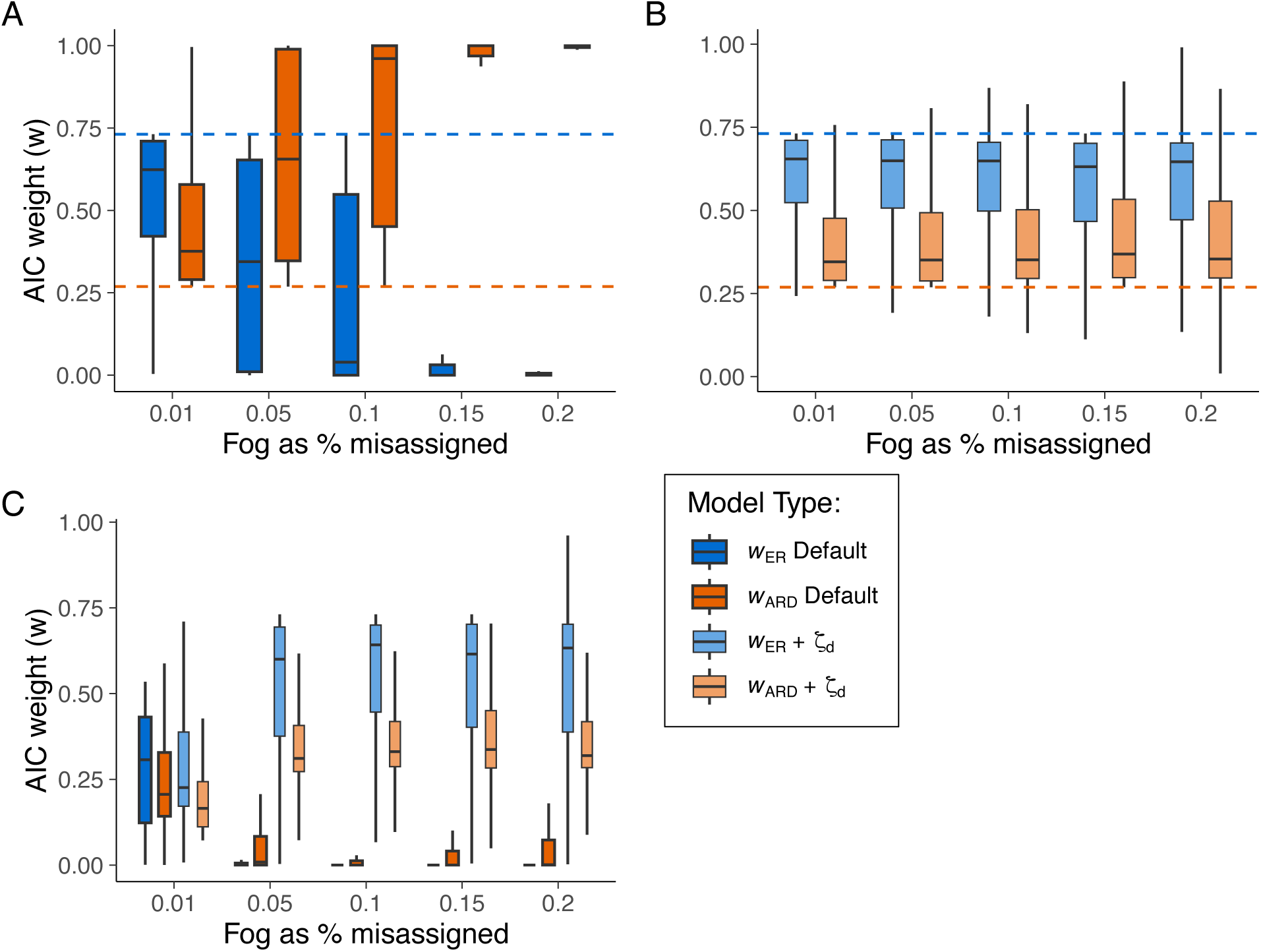
Summaries of model support based on Akaike weight (*w*) for equal rates (ER) and all rates different (ARD) continuous-time Markov models fit (**A**) without tip fog and (**B**) fit including tip fog as a parameter, or (**C**) pooled together as part of an inclusive model set. The generating model for these simulations was an equal rates (ER) continuous-time Markov model (*q01* = *q10* = 0.025 transitions Myr^-1^) with increasing levels of tip fog. To simulate tip fog, we randomly altered the observed state of 1%, 5%, 10%, 15%, or 20% of taxa to be the reverse of its true state. Dashed lines in (**A**) and (**B**) represent the null expectation of the Akaike weight as the average Akaike weight if we assume an equal likelihood across all models.

To illustrate this further, we also included an HMM model that contained two distinct rate classes (*q01A* = *q10A* and *q01B* = *q10B*) and an additional transition rate governing the transition among them (i.e., three transition rates total). When this model was included as part of the default model set, the ARD and HMM models competed for the highest Akaike weight, with the ER model showing very little support generally for tip fog values >1% (Fig. S4; Table S4). Interestingly, the HMM consistently had the highest support in trees with clade ages >10 Myr (Table S4). When the tip fog probability was estimated as part of the model, support for the HMM consistently converged toward the null Akaike weight, which was substantially lower than either the ER or ARD models (Fig. S4).

We were also curious about the performance of the +ζ_*d*_ models when included as part of a broader set that also included models that did not estimate ζ_*d*_. It may be that even though the ER + ζ_*d*_fits well when compared to other +ζ_*d*_ models, the added complexity of the additional parameter reduces the power to detect tip fog when it is present. However, when we pooled the default and tip fog model sets together and recalculated the Akaike weights, the ER + ζ_*d*_ model emerged as the best model across all fog values except ζ_*d*_ = 1% (Fig. 4C; Table S3). In that case, these results suggest that such low fog values are often difficult to distinguish from no fog, which is expected.

### Extending to state-speciation and extinction models

The approach for estimating tip fog from discrete traits extends naturally to state-speciation and extinction models (SSE; Maddison et al., 2007; FitzJohn et al., 2009; Beaulieu & O’Meara, 2016). We parameterize the initial conditions for *DN,i*(*t*) – the probability that a lineage observed in state *i* at time *t* – to be 1 – ζ*_i,j_* for the observed state and ζ*_i,j_* for the alternative state. The initial conditions for *Ei*(*t*), the probability that a lineage in state *i* at time *t* would go completely extinct by the present, remains unaltered. The likelihood calculation then proceeds down the tree as usual.

When the input tree represents a sample of all extant species within a focal clade, the initial conditions for *DN,i*(*t*) must also account for the sampling fraction, *fi*, which specifies the probability that a species with true state *i* is sampled and included in the tree. For example, if a tip is observed in state *0* at the present, the initial conditions for *DN,0*(*t*=0) = *f0* (1 - ζ_*d*_) and *DN,1*(*t*=0) = *f1* ζ_*d*_ for the alternative state. The initial probability for *E0*(0) remains the probability of a lineage not being present in the phylogeny, either by going extinct or not being sampled, and is therefore set as *Ei*(*t*=0) = 1 − *f0*. These modifications have been implemented in the hidden state-speciation and extinction model in the R package *hisse* (Beaulieu & O’Meara 2016).

Unincorporated tip fog should have two effects on the estimation and interpretation of parameters in an SSE model. First, tip fog should erroneously inflate transition rates, though not to the degree that they are in continuous-time Markov models. This is because SSE models jointly estimate speciation, extinction, and state transition processes, making it less likely for the model to attribute too much to transition rates alone. Instead, character misassignments are more likely to be absorbed as part of the variation in speciation and/or extinction rates. However, this benefit comes with a trade-off: the model may homogenize diversification rates among observed states, which can lead to increased support for models that assume some form of character-independence. In other words, some of the tips observed in state *0* are actually in state *1*, and vice versa, making the states seem more similar from a diversification standpoint than they should be.

To investigate how SSE models behave with tip fog, we simulated scenarios where the turnover rate (λ + μ) for state *1* was nearly double that of state *0*. Specifically, we simulated 100 trees with λ_0_ = 0.22 events Myr^-1^ and λ_1_ = 0.42 events Myr^-1^, with extinction rates set to 75% of the speciation rates in both cases, and equal transition rates of 0.025 transitions Myr^-1^. Each tree started in state *0* and evolved for 50 Myr, resulting in an average of about 200 taxa per tree. To avoid patterns from simulation time bias or inflating the effect of tip fog by stopping at a fixed number of taxa, which can result in a clade with zero length branches, we terminated simulations at a pre-specified time. Tip fog was introduced by randomly altering the true state of 1%, 5%, 10%, 15%, or 20% of taxa to the incorrect state. For each simulation replicate, we fit two sets of six models, including both character-independent (e.g., CID-2) and character-dependent models (e.g., BiSSE), with either equal or asymmetric transition rates (see Table S4). Each model set either ignored tip fog (referred to as “default”) or estimated its probability (+ζ_*d*_). We calculated a weighted harmonic average of the transition rates across model fits using Akaike weights. Estimates of turnover rates were summarized as a weighted harmonic mean of the rates represented at the tips of the tree. That is, for each model, the marginal probability of each state (and rate class for CID-2) was computed for every tip, and then computed as the weighted harmonic mean across all models using the Akaike weights.

As expected, our simulations revealed that transition rates became increasingly inflated with higher degrees of tip fog when not accounting for tip fog in the model (Fig. 5A). In addition, as expected the magnitude of the rate inflation is muted compared to the continuous-time Markov models for the same tip fog (Fig. 3). In contrast, when tip fog was estimated as part of the model, the individual transition rates generally remained close to their true values as did the estimates of the tip fog probability (Fig. 5C).

**Figure 5.**
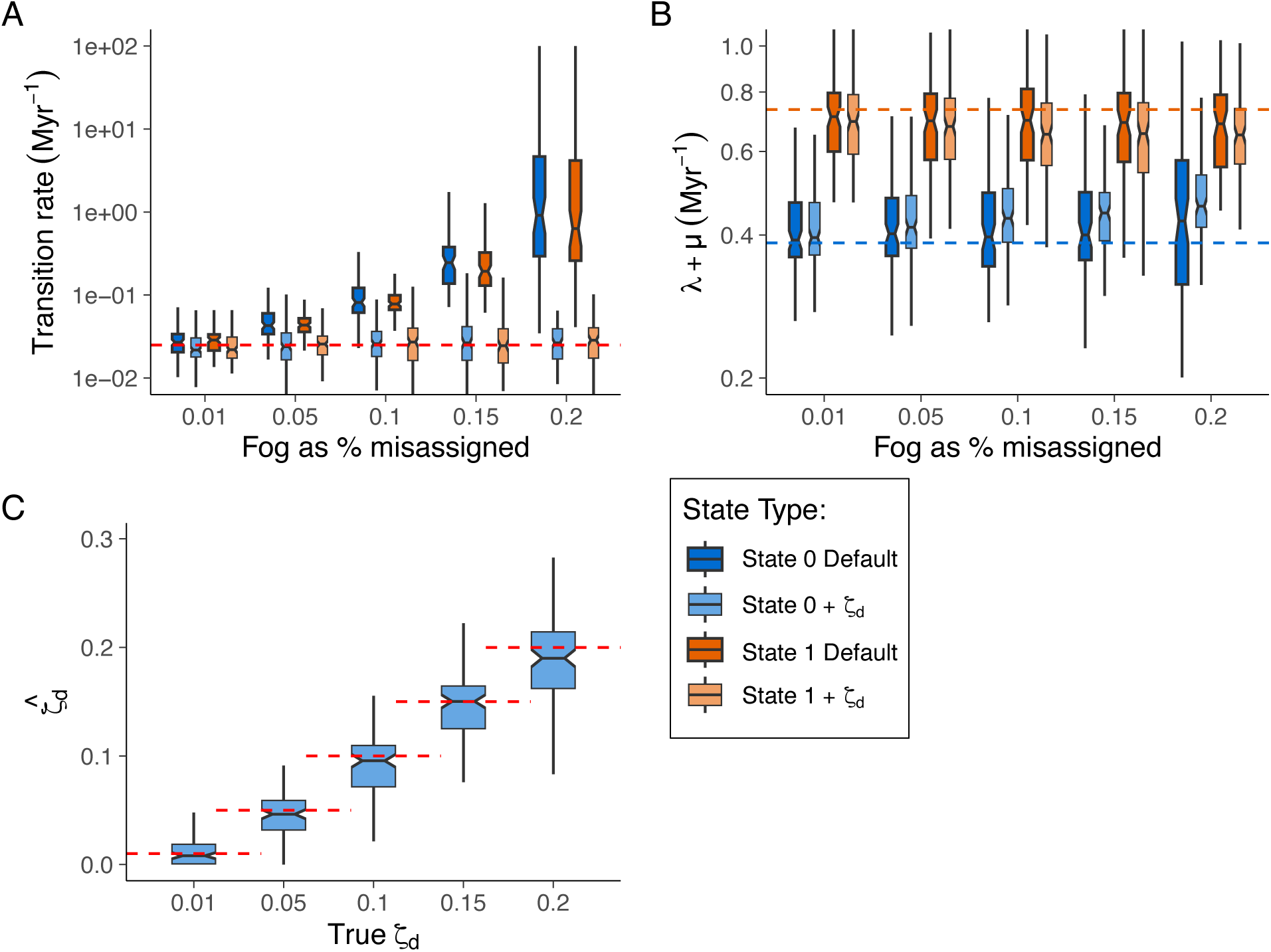
Uncertainty in estimating (**A**) transition rates and (**B**) turnover rates under a state-speciation and extinction model with increasing levels of tip fog. The generating model was a character-dependent model (CD), where state *1* to have turnover rate (λ_1_ + μ_1_ = 0.735 events Myr^-1^) that was nearly 2x the rate of state *0* (λ_0_ + μ_0_ = 0.385 events Myr^-1^), with state transitions between *0* and *1* set at 0.025 transitions Myr^-1^; extinction fraction was set at 0.75 for both regimes. To simulate tip fog, we randomly altered the observed state of 1%, 5%, 10%, 15%, or 20% of taxa to be the reverse of its true state. Data sets were evaluated two sets of six models, including both character-independent (e.g., CID-2) and character-dependent models (e.g., BiSSE), with either equal or asymmetric transition rates (see Table S3) Each model set either ignored tip fog (Default) or estimated its probability (+ζ_*d*_). Rate estimates within each of the model classes were then summarized by calculating a weighted harmonic mean of each rate parameter using the Akaike weights. Darker boxes indicate transition rates summarized across models excluding tip fog (Default); the less saturated boxes represent transition rates summarized across models that estimated tip fog (+ζ_*d*_). Dashed blue and orange lines indicate the generating values for state *0* and state *1*, respectively. (**C**) depicts uncertainty in ζ_*d*_ estimates, with the dashed red line indicating the true ζ_*d*_.

Unexpectedly, turnover rates tended to converge to a single intermediate value as the degree of tip fog increased, regardless of whether tip fog was estimated (Fig. 5B). We suspect this pattern arises for two distinct reasons. First, in the default model set that does not include ζ_*d*_, the convergence appears to be driven by an increasing model weight toward character-independent models as the level of tip fog increases (Fig. 6A, Table S5). There is increased support for models that do not differentiate based on the observed character states, leading to more uniform estimates of turnover rates across different states. In contrast, for the +ζ_*d*_models, support for character-dependent models remained stable across different levels of simulated tip fog (Fig. 6B).

**Figure 6.**
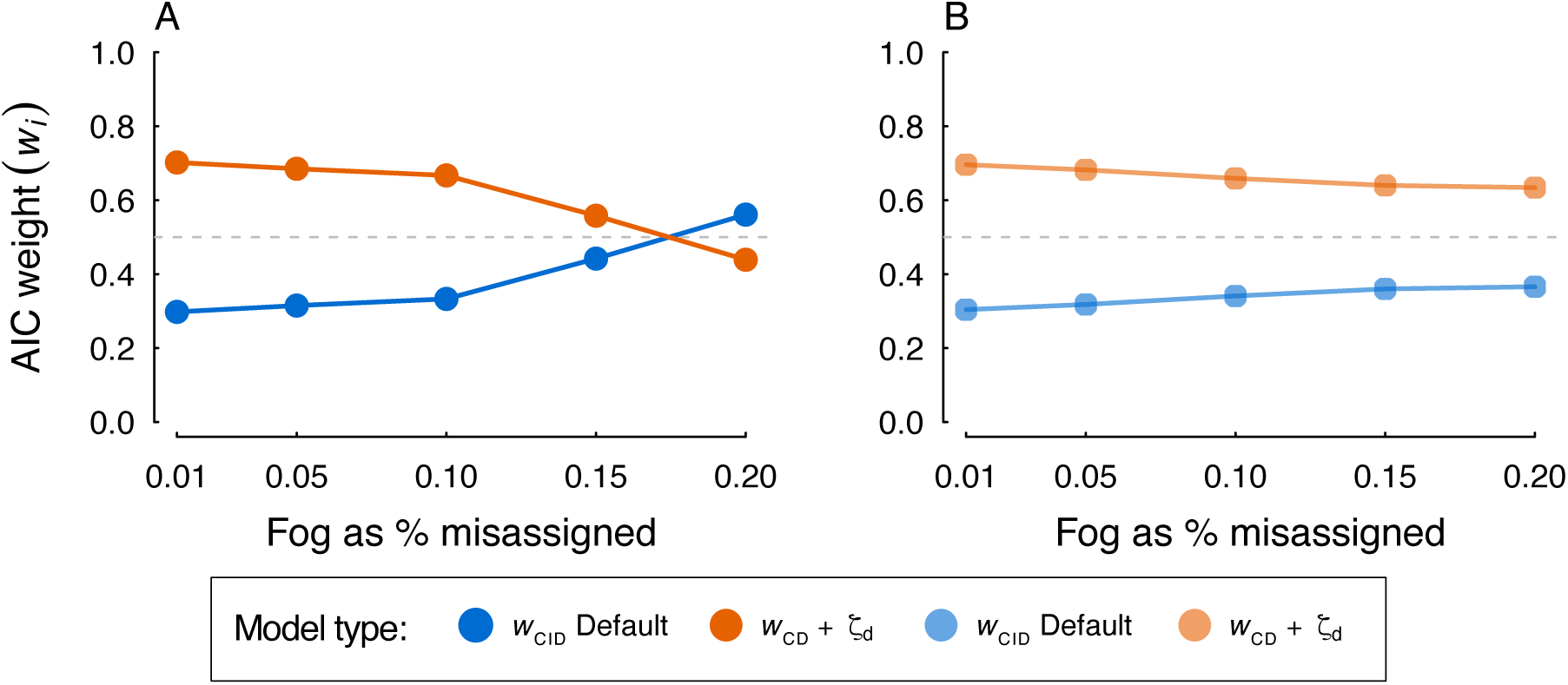
Summary of the cumulative support for character-dependence (CD; turnover rate depends on the state) and character-independence (CID; turnover rates do not depend on the state) when the generating model is a character-dependent model that contains increasing levels of tip fog. Panel (**A**) shows how support for character-independence increases with increasing tip fog when tip fog is ignored (Default), whereas (**B**) when tip fog is estimated as part of the model (+ζ_*d*_) support for character-dependent models remains stable across different levels of simulated tip fog.

The convergence of turnover rates despite the stability in model support for character-dependence within the +ζ_*d*_ models is likely due to how tip rates are summarized when tip fog is present. Consider a BiSSE model where the turnover rate is 0.35 events Myr^-1^ for state *0* and 0.70 events Myr^-1^ for state *1*. In the absence of tip fog, the rate for a given tip would simply be the observed state since there is no uncertainty as to the true state. However, with the tip fog probability being estimated a tip might be observed in say state *1*, but there is some uncertainty as to whether it is state *0* instead. Thus, we must account for this when summarizing the rates within a given model when tip fog probability is included, and this tends to homogenize diversification rates as tip fog increases. For instance, if a model is estimated to have a 5% tip fog probability, the tip rate for a taxon observed in state *1* for that model is calculated as a weighted average such that (0.35 x 0.05) + (0.70 x 0.95) = 0.68 events Myr^-1^. Now, if the tip fog probability increases to, say, 20%, as we did in our simulations, the tip rate would adjust downwards to (0.35 x 0.2) + (0.70 x 0.8) = 0.63 events Myr^-1^. This adjustment would also cause the turnover rate for tips observed in state *0* to gradually increase, contributing to the overall homogenization of turnover rates, even with clear support for the character-dependent models included in the model set.

## Discussion

While the importance of tip fog in continuous traits has long been acknowledged, we were surprised by how accurately it can be estimated from both continuous and discrete data directly. This is significant because tip fog is not just simply adding another parameter — it represents the extent to which tip data distorts or misrepresents the underlying reality. Given the feasibility of estimating it and the detrimental effects of setting it to zero, we have updated several popular software packages to estimate tip fog, and we recommend others do the same. Anyone converting biological variation into discrete data knows this inevitably leads to fuzzy cases, and this fuzziness is even more pronounced with continuous variation. Ignoring tip fog can result in confidently incorrect conclusions rather than mere uncertainty. Even if practitioners are hesitant to increase model complexity by estimating an additional parameter, selecting an arbitrary value of say, 10%, is likely more accurate than the current practice of assuming tip fog is 0% (but see Kuhner & McGill (2014) about the risks of incorrectly assuming too much fog).

Our implementation of tip fog is straightforward. For continuous traits, we use a constant value across all tips, while for discrete traits, we follow a similar approach in our simulations, though we also allow for varying error rates depending on observed states. For example, in traits like parental care, it is more likely that species with this trait might be missed rather than incorrectly reported as having it. However, there are opportunities to increase complexity. Tip fog for continuous traits could be modeled as a proportion of the observed state instead of a fixed standard deviation. Moreover, different amounts of fog could be estimated for species categorized by factors such as observations from herbaria versus field studies, species with extensive records versus those with fewer, and observations made by undergraduates versus faculty. Incorporating the number of observations per species could also refine estimates of continuous tip fog. Exploring different regimes of tip fog across the phylogenetic tree offers additional avenues for improvement (*sensu* Ives et al., 2007).

We hasten to acknowledge that tip fog is not the only source of uncertainty in macroevolutionary studies. Uncertainty in topology and branch lengths, as well as unincorporated heterogeneity in the evolutionary process, also likely play significant roles. Properly incorporating tip fog does not negate the need to consider these other factors. However, just as incorporating tree uncertainty by conducting analyses across a set of trees is essential, so is incorporating tip fog.

The concept of tip fog presents an opportunity to revisit the basic principles of model selection. Models with or without tip fog estimation can be compared using metrics such as AIC, AICc, or BIC, all of which account for model complexity. The model with the lowest score on these metrics offers the best balance between complexity and fit. For instance, if a model that includes tip fog estimation has a ΔAIC of 0, and a model that forces tip fog to zero has a ΔAIC of 1.4, the model estimating tip fog is superior. However, it is not uncommon to encounter model choice being based on requiring that a more complex model outperform a simpler one by a certain arbitrary margin before considering it (i.e., ΔAIC > 2). However, such an approach is neither necessary nor appropriate (Burnham & Anderson, 2004).

We also note that tip fog is distinct from approaches that account for polymorphism in tip data, although they are related. For instance, consider a character with states yellow, white, and red, where one species exhibits polymorphism with some flowers being yellow and others white, while most species in the clade are uniformly one of the three colors. In such a case, the likelihood calculation would start with *P*(*Oi* = yellow) = *P*(*Oi* = white) = 1 for that species [rather than 0.5 for each, as noted by Felsenstein (2004)]. This differs from errors such as a researcher misassigning a yellow specimen as white due to poor lighting. Despite this distinction, tip fog can still be applied alongside polymorphism to improve model accuracy. In fact, this might be particularly useful for models of biogeography where polymorphic scoring is a central feature (Ree & Smith 2008; Bätscher & de Vos, 2024).

Our study focuses on tip fog within traditional macroevolutionary models, which typically analyze one or a few characters. However, tip fog can significantly impact models that handle multiple characters, such as those used for inferring phylogenetic trees and networks (e.g., Kuhner & McGill). One major component of tip fog is sequencing error, which is likely more substantial than is typically acknowledged. Incorporating tip fog as a default option in tree inference is particularly sensible given the large volumes of data often available. Yet, popular tree inference programs like RAxML-NG (Kozlov et al., 2019) and IQ-TREE (Minh et al., 2020) currently lack this capability. The omission of tip fog is especially critical for branch length estimates, as unaccounted-for tip fog tends to lengthen terminal branches (which make up over half of a tree’s edges). This can lead to an inaccurate estimation of tree age if a rate calibration is used. When using multiple fossils or other calibrations, the effect on overall tree age is less predictable but generally results in an increased ratio of terminal to internal branch lengths. In any event, while some models for cancer tumor phylogenies incorporate error expectations (Davis & Navin 2016), sequencing error remains a significant concern in traditional phylogenetic studies (see also Ho et al., 2005).

We note several important caveats. Our analysis has focused on predictable errors, such as a species mean being off by 10% or a 20% chance of misassigning a species’ state as woody instead of herbaceous. However, we have not addressed more extraordinary sources of error, such as entering a unitless mass in milligrams for one species while using grams for others, confusing range data between a plant and an insect due to homonyms, omitting the sign for longitude, or recording a missing value as −99 and then incorrectly treating this as a valid measurement. Tip fog, as we incorporate it, is unlikely to address these types of errors effectively. While tip fog can account for certain uncertainties, it relies on the data being fundamentally accurate. Additionally, tip fog does not correct errors in tree or network topology or branch lengths – these also remain important to incorporate. While our work uses likelihood and AIC, we expect similar results using Bayesian methods or using model selection criteria beyond AIC. Even with Bayesian methods, which deal quite well in uncertainty, existing approaches essentially put full prior weight on the data being completely right, not allowing any exploration about the possibility of nonzero tip fog (but see Revell & Reynolds, 2012). Either adding one or more tip fog parameters or allowing a looser coupling between observed and actual states, would allow Bayesian methods to incorporate this important factor.

Finally, several questions remain unanswered. For instance, empirical estimates of variability at tips, such as the standard deviation of samples, might underrepresent tip fog because they do not account for factors affecting all modern samples, such as environmental plasticity. This issue has not been explored in our study; it remains uncertain whether estimating tip fog directly is more effective than relying on empirical estimates of tip variability or if adding an “additional” fog parameter to the model would be beneficial. While we have incorporated tip fog into SSE models for discrete traits, we have not applied it to SSE models of continuous data (e.g., QuaSSE; FitzJohn, 2010). We also have not explored multivariate models, as discussed by Felsenstein (2008). Additionally, some simulation results were unexpected. For example, we anticipated that tip fog would significantly improve estimates of θ in an Ornstein-Uhlenbeck model, but the effect appears less pronounced than expected. The presence and impact of tip fog in empirical studies are still largely unexplored (but see some examples in the Supplementary Materials); analyzing even more existing datasets to estimate tip fog and assess the consequences of ignoring it would be a valuable next step.

### Conclusions

Melville warned that those who seek to understand whales must risk their boats being crushed. Similarly, many comparative analyses are at risk of failing due to unrecognized variation from a myriad of sources — what we term “tip fog.” However, this risk can be mitigated by incorporating tip fog into our standard models, which will improve the accuracy of our inferences and avoid the pitfalls of confidently incorrect conclusions. As we navigate the complexities of biological data, making tip fog a standard consideration will provide more reliability to our analyses.

## Acknowledgments

Conversations with Andy Alverson, James Boyko, James Pease, Tomomi Parins-Fukuchi, Nat Walker-Hale, Jakob Berv, and Stephen Smith were invaluable providing constructive feedback. We are especially grateful to Josef Uyeda for his insightful discussions on the concept of tip fog and for pointing out the extremity of our original simulations, which greatly improved our approach.

## Notes

**Funding:** This study was funded by grants from the National Science Foundation (grant DEB−1916558 awarded to J.M.B. and grant DEB−1916539 awarded to B.C.O.).

https://github.com/thej022214/MeasurementError

https://zenodo.org/doi/10.5281/zenodo.13685561

### Competing Interest Statement

The authors have declared no competing interest.

### Summary of Updates

A reader pointed out that our original simulations for the continuous trait portion of the manuscript were quite extreme. We agree, and decided to redo this portion of the analyses with more reasonable tip fog values. We also were urged to demonstrate empirical estimates for tip fog in this part of the part, which we've added as well. All interpretations and conclusions remain unchanged.

## References

Alencar, L.R.V., Schwery, O., Gade, M. R., Domínguez-Guerrero, S. F., Tarimo, E, Bodensteiner, B. L., Uyeda, J.C., & Muñoz, M. M. (2024). Opportunity begets opportunity to drive macroevolutionary dynamics of a diverse lizard radiation. Evolution Letters, In press, doi: 10.1093/evlett/qrae022.

Bätscher, L., & de Vos, J. M. (2024). Avoiding impacts of phylogenetic tip-state-errors on dispersal and extirpation rates in alpine plant biogeography. Journal of Biogeography, 51(6), 1104–1116. doi: 10.1111/jbi.14811

Beaulieu, J. M. & O’Meara, B. C. (2016). Detecting hidden diversification shifts in models of trait-dependent speciation and extinction. Systematic Biology, 65(4), 583–601. doi: 10.1093/sysbio/syw022

Beaulieu, J. M., Jhwueng, D.-C., Boettiger, C., & O’Meara, B. C. (2012). Modeling stabilizing selection: expanding the Ornstein-Uhlenbeck model of adaptive evolution. Evolution, 66(8), 2369–2383. doi: 10.1111/j.1558-5646.2012.01619.x

Beaulieu, J. M., O’Meara, B. C., & Donoghue, M. J. (2013). Identifying hidden rate changes in the evolution of a binary morphological character: the evolution of plant habit in campanulid angiosperms. Systematic Biology, 62(5), 725–737. doi: 10.1093/sysbio/syt034

Boyko, J. D., & Beaulieu, J. M. (2021). Generalized hidden Markov models for comparative datasets. Methods in Ecology and Evolution, 12(3), 468-478. doi: 10.1111/2041-210X.13534

Boyko, J. D., & O’Meara, B. C. (2024). dentist: quantifying uncertainty by sampling points around maximum likelihood estimates. Methods in Ecology and Evolution, 15(4), 628–638. doi: 10.1111/2041-210X.14297

Burnham, K. P., & Anderson, D. R. (2002). Model selection and multimodal inference. Springer Press.

Butler, M. A., & King, A. A. (2004). Phylogenetic comparative analysis: a modeling approach for adaptive evolution. The American Naturalist, 164(6), 683–695. doi: 10.1086/426002

Cheverud, J. M., Dow, M. M., & Leutenegger, W. (1985) The quantitative assessment of phylogenetic constraints in comparative analyses: sexual dimorphism in body weight among primates. Evolution, 39(6), 1335–1351. doi: 10.1111/j.1558-5646.1985.tb05699.x

Davis, A. & Navin, N. E. (2016). Computing tumor trees from single cells. Genome Biology, 17, 113. doi: 10.1186/s13059-016-0987-z

Felsenstein, J. (2004). Inferring phylogenies. Sinauer Associates.

Felsenstein, J. (2008). Comparative methods with sampling error and within-species variation: contrasts revisited and revised. The American Naturalist, 171(6), 713–725. doi: 10.1086/587525

FitzJohn, R. G. (2010). Quantitative traits and diversification. Systematic Biology, 59(6), 619–633. doi: 10.1093/sysbio/syq053/

FitzJohn, R. G., Maddison, W. P., & Otto, S. P. (2009). Estimating trait-dependent speciation and extinction rates from incompletely resolved phylogenies. Systematic Biology, 58(6), 595–611. doi: 10.1093/sysbio/syp067

Goolsby, E. W., Bruggeman, J., & Ané, C. (2016). Rphylopars: fast multivariate phylogenetic comparative methods for missing data and within-species variation. Methods in Ecology and Evolution, 8(1), 22–27. doi: 10.1111/2041-210X.12612

Han, M. V., Thomas, W. C., Lugo-Martinez, J., & Hahn, M. W. (2013). Estimating gene gain and loss rates in the presence of error in genome assembly and annoting using CAFE 3. Molecular Biology and Evolution, 30(8), 1987–1997. doi: 10.1093/molbev/mst100.

Hansen, T. F. & Bartoszek, K. (2012). Interpreting the evolutionary regression: the interplay between observational and biological errors in phylogenetic comparative studies. Systematic Biology, 61(3), 413–425. doi: 10.1093/sysbio/syr122.

Harmon, L. J. & Losos, J. B. (2005). The effect of intraspecific sample size on type I and type II error rates in comparative studies. Evolution, 59(12), 2705–2710. doi: 10.1111/j.0014-3820.2005.tb00981.x

Ho, S. Y. W., Phillips, M. J., Cooper, A., & Drummond, A. J. (2005). Time dependency of molecular rate estimates and systematic overestimation of recent divergence times. Molecular Biology and Evolution, 22(7), 1561–1568. doi: 10.1093/molbev/msi145

Ho, S.Y.W., Lancer, R., Phillips, M. J., Barnes, I., Thomas, J. A., Kolokotronis, S.-O., & Shapiro, B. (2007). Systematic Biology, 60(3), 366–375. doi: 10.1093/sysbio/syq099

Höhna, S., Landis, M. J., Heath, T. A., Boussau, B., Lartillot, N., Moore, B. R., Huelsenbeck, J. P., & Ronquist, F. (2016). RevBayes: Bayesian phylogenetic inference using graphical models and an interactive model-specification language. Systematic Biology, 65(4), 726–736. doi: 10.1093/sysbio/syw021

Ingram, T., & Mahler, D. L. (2013). SURFACE: detecting convergent evolution from comparative data by fitting Ornstein-Uhlenbeck models with stepwise Akaike Information Criterion. Methods in Ecology and Evolution, 4(5), 416–425. doi: 10.1111/2041-210X.12034

Ives, A. R., Midford, P. E., & Garland, T., Jr. (2007). Within-species variation and measurement error in phylogenetic comparative methods. Systematic Biology, 56(2), 252–270. doi: 10.1080/10635150701313830

Jackson, C. H., Sharples, L. D., Thompson, S. G., Duffy, S. W., & Couto, E. (2003). Multistate Markov models for disease progression with classification error. Journal of the Royal Statistical Society Series D: The Statistician, 52(2), 193–209. doi: 10.1111/1467-9884.00351

Kozlov, A. M., Darriba, D., Flouri, T., Morel, B., & Stamatakis, A. (2019). RAxML-NG: a fast, scalable and user-friendly tool for maximum likelihood of phylogenetic inference. Bioinformatics, 35(21), 4453–4455. doi: 10.1093/bioinformatics/btz305

Kuhner, M. K., & McGill, J. (2014). Correcting for sequencing error in maximum likelihood phylogeny inference. G3 (Bethesda), 4(12), 2545-2552. doi: 10.1534/g3.114.014365

Labra, A., Pienaar, J., & Hansen, T. F. (2009). Evolution of thermal physiology in Liolaemus lizards: adaptation, phylogenetic inertia, and niche tracking. The American Naturalist, 174(2), 204–220. doi: 10.1086/600088

Lynch, M. (1991). Methods for the analysis of comparative data in evolutionary biology. Evolution, 45(5), 1065–1080. doi: 10.1111/j.1558-5646.1991.tb04375.x

Maddison, W. P., Midford, P. E., & Otto, S. P. (2007). Estimating a binary character’s effect on speciation and extinction. Systematic Biology, 56(5), 701–710. doi: 10.1080/10635150701607033

Minh, B. Q., Schmidt, H. A., Chernomor, O., Schrempf, D., Woodhams, M. D., von Haeseler, A., & Lanfear, R. (2020). IQ-TREE 2: new models and efficient methods for phylogenetic inference in the genomic era. Molecular Biology and Evolution, 37(5), 1530–1534. doi: 10.1093/molbev/msaa015

O’Meara, B. C., & Beaulieu, J. M. (2024). Noise leads to the perceived increase in evolutionary rates over short time scales. PLoS Computational Biology, Accepted.

O’Meara, B. C., Ané, C., Sanderson, M. J., & Wainwright, P. C. (2006). Testing for different rates of continuous trait evolution using likelihood. Evolution, 60(5), 922–933. doi: 10.1111/j.0014-3820.2006.tb01171.x

Pennell, M. W., Eastman, J. M., Slater, G. J., Brown, J. W., Uyeda, J. C., FitzJohn, R. G., Alfaro, M. E., & Harmon, L. J. (2014). geiger v2.0: an expanded suite of methods for fitting macroevolutionary models to phylogenetic trees. Bioinformatics, 30(15), 2216–2218. doi: 10.1093/bioinformatics/btu181

Rambaut, A., Ho, S. Y. W., Drummond, A. J., & Shapiro, B. (2008). Accommodating the effect of ancient DNA damage on inferences of demographic histories. Molecular Biology and Evolution, 26(2), 245–248. doi: 10.1093/molbev/msn256

Ree, R. H., & Smith, S. A. (2008). Maximum likelihood inference of geographic range evolution by dispersal, local extinction, and cladogenesis. Systematic Biology, 57(1), 4–14. doi: 10.1080/10635150701883881

Revell, L. J., & Reynolds, G. (2012). A new Bayesian method for fitting evolutionary models to comparative data with intraspecific variation. Evolution, 66(9), 2697–2707. doi: 10.5061/dryad.7fv08k72

Stadler, T. (2011). Simulating trees with a fixed number of extant species. Systematic Biology, 60(5), 676–684. doi: 10.1093/sysbio/syr029

Sylvestro, D., Kostikva, A., Litsios, G., Pearman, P. B., & Salamin, N. (2015). Measurement errors should always be incorporated in phylogenetic comparative analysis. Methods in Evolution and Evolution, 6(3), 340–346. doi: 10.1111/2041-210X.12337

Uyeda, J. C., & Harmon, L. J. (2014). A novel Bayesian method for inferring and interpreting the dynamics of adaptive landscapes from phylogenetic comparative data. Systematic Biology, 63(6), 902–918. doi: 10.1093/sysbio/syu057

