## Supplemental Materials for "Navigating “tip fog”: Embracing uncertainty in tip measurements"

#### TABLE OF CONTENTS


A. **Supplementary Figures**

B. **Supplementary Tables**

C. **Empirical tip fog estimates**

D. **References**

### A. Supplementary Figures

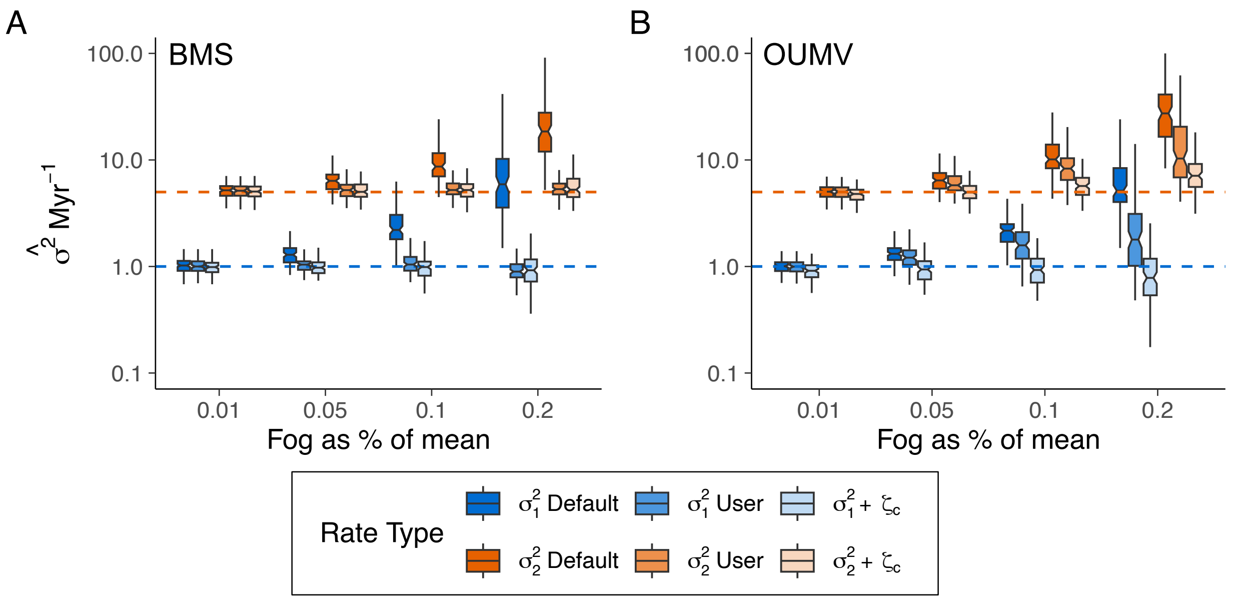


**Figure S1.** Uncertainty in estimating the evolutionary rate ($\sigma^2$) as a function of tip fog, with the generating model assuming various (**A**) multiple-rate Brownian motion (BMS) and (**B**) multiple mean, multiple-rate Ornstein-Uhlenbeck (OUMV) models. The generating model in both panels is a rate for regime 2 that is 5-times that of regime 1. Tip fog was simulated by resampling each tip value from a normal distribution centered at the individual species mean and with a standard deviation that was a percentage of the mean. Data sets were then evaluated under BM1, BMS, OU1, OUM, and OUMV models, with rates summarized using a weighted harmonic mean based on Akaike weights (see text). Darker boxes indicate rate summarized across models excluding tip fog (Default), which show an upward bias in evolutionary rates as fog levels increase, regardless of the regime. The less saturated boxes represent rates summarized across models that either calculate tip fog from simulated "measurements" (+ User) or estimate tip fog (+$\zeta_c$), where evolutionary rates generally align more closely with true values for BMS, but for OUMV the +User estimates are upward biased. Dashed blue and orange lines indicate the generating values for regimes 1 and 2, respectively.

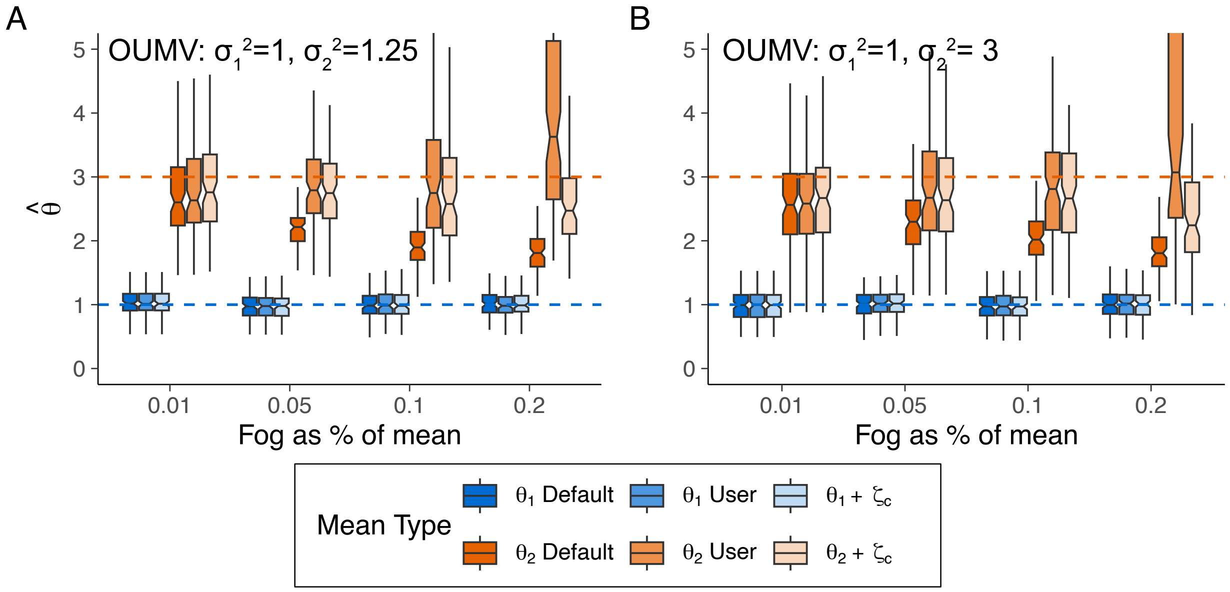


**Figure S2.** Uncertainty in estimating the trait means ($\theta_i$) when the generating model is a multiple-mean, multiple-rate Ornstein-Uhlenbeck model (OUMV), and the simulated data sets contained differing levels of tip fog. In both cases, the generating model assumes that the trait mean for regime 2 was 3x that of regime 1. Panel (**A**) depicts a scenario where the rate for regime 2 has a rate that is 1.25 times that of regime 1, whereas (**B**) depicts a scenario where regime 2 has a rate that is 3 times that of regime 1. Data sets were evaluated under BM1, BMS, OU1, OUM, and OUMV models, with rates summarized using a weighted mean based on Akaike weights (see text). Darker boxes indicate $\theta_i$ summarized across models excluding tip fog (Default); the less saturated boxes represent $\theta_i$ summarized across models that that either calculate tip fog from simulated "measurements" (+ User) or estimated tip fog ($+\zeta_c$). Dashed blue and orange lines indicate the generating values for regimes 1 and 2, respectively. In both scenarios, as the amount of tip fog increases, estimates for $\theta_2$ are increasingly underestimated, irrespective of whether tip fog was estimated.

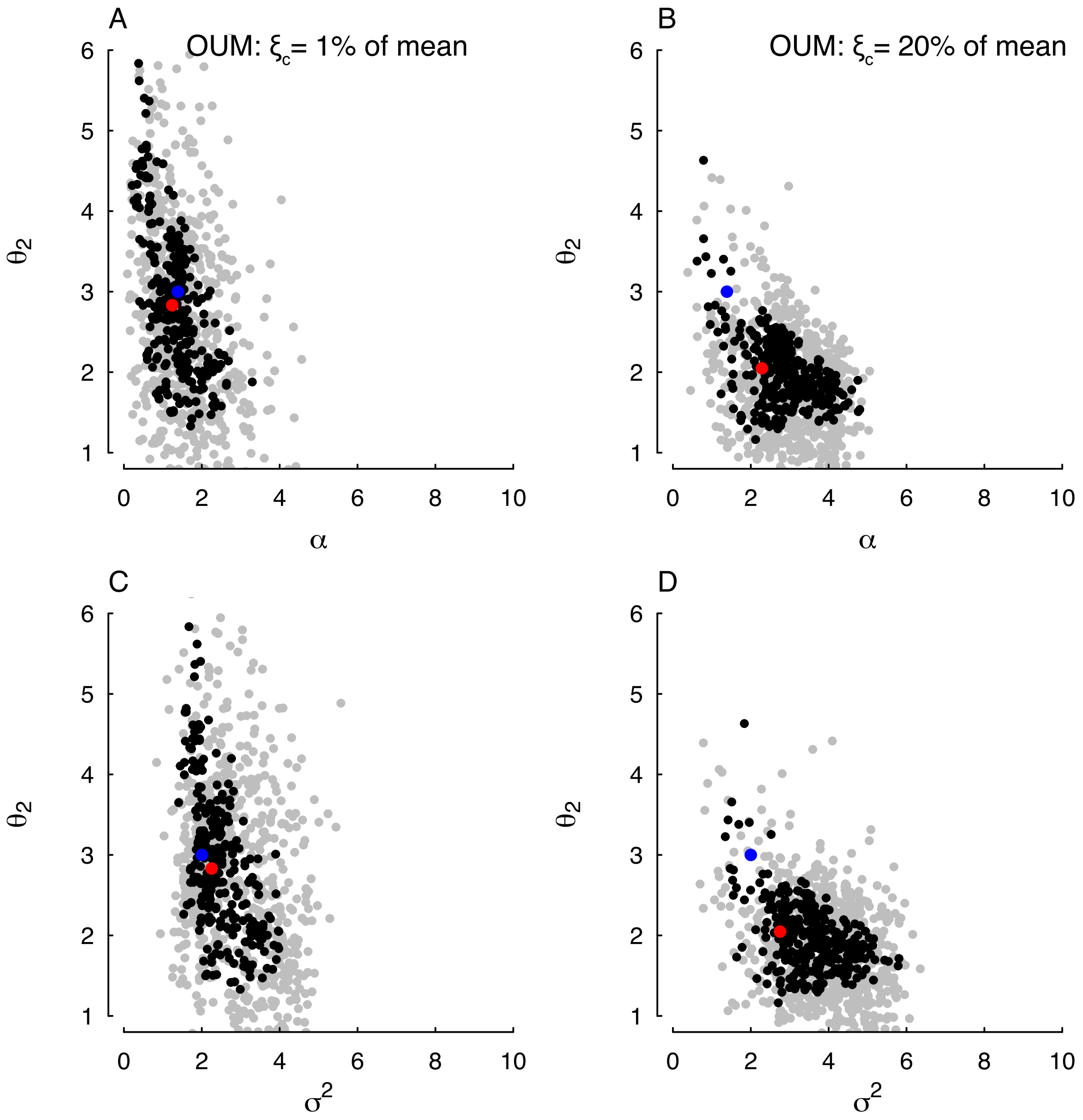


**Figure S3.** Bivariate plots, with sampled parameter values represented by dots, with gray dots indicating values outside the confidence region and black dots inside. In an effective analysis, the black region in these plots should form an ellipsoid shape, surrounded by gray. To obtain a conservative estimate from the points, such as those in Fig. 1 and 2, a rectangular prism was placed around the clusters of black points. The range of values, such as the proportion of hyperbolic weight, was derived from calculations at the vertices of this prism. The overall approach of dentist (Boyko & O'Meara, 2024) resembles Markov Chain Monte Carlo in Bayesian analysis but does not rely on prior distributions. Instead, it focuses on establishing bounds rather than defining a region that comprises a certain proportion of overall probability. Examining the confidence regions surrounding estimates of $\theta_2$, $\alpha$, and $\sigma^2$ using reveals that tip fog introduces greater uncertainty in estimates of $\alpha$ and $\sigma^2$, which incidentally impacts estimates of $\theta_2$, even when $\zeta_c$ is included in the model.

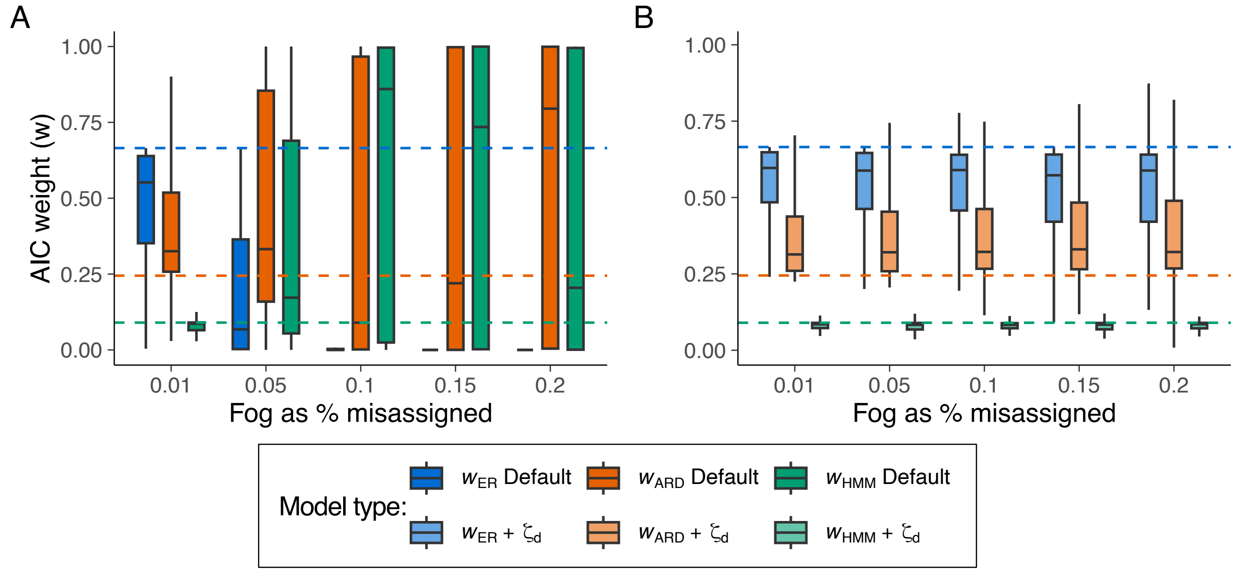


**Figure S4.** Summaries of model support based on Akaike weight (*w*) for equal rates (ER) and all rates different (ARD) continuous-time Markov models fit, and an HMM model (see text) (**A**) without tip fog and (**B**) fit including tip fog as a parameter. The generating model for these simulations was an equal rates (ER) continuous-time Markov model ($q_{01} = q_{10} = 0.025$ transitions $Myr^{-1}$) with increasing levels of tip fog. To simulate tip fog, we randomly altered the observed state of 1%, 5%, 10%, 15%, or 20% of taxa to be the reverse of its true state. Dashed lines in (**A**) and (**B**) represent the null expectation of the Akaike weight as the average Akaike weight if we assume an equal likelihood across all models.

### A. Supplemental Tables

**Table S1.** Summary of model support for simulated scenarios of continuous-trait evolution that contained increasing levels of tip fog, assessed by calculating the average Akaike weight (w) for all models assessed the fit.

| | Default | | | | | + User-supplied $\xi_c$ | | | | | + $\xi_c$ | | | | |
|---|---|---|---|---|---|---|---|---|---|---|---|---|---|---|---|
| Model | $BM1_w$ | $BMS_w$ | $OU1_w$ | $OUM_w$ | $OUMV_w$ | $BM1_w$ | $BMS_w$ | $OU1_w$ | $OUM_w$ | $OUMV_w$ | $BM1_w$ | $BMS_w$ | $OU1_w$ | $OUM_w$ | $OUMV_w$ |
| Half-life = ∞; $\sigma_1^2 = 1, \sigma_2^2 = 1.25$; $\theta_0 = 1$ | | | | | | | | | | | | | | | |
| $\xi_c$=1% | 0.264 | 0.285 | 0.193 | 0.136 | 0.122 | 0.277 | 0.286 | 0.188 | 0.132 | 0.117 | 0.293 | 0.299 | 0.155 | 0.133 | 0.119 |
| 2.5% | 0.196 | 0.273 | 0.211 | 0.144 | 0.176 | 0.282 | 0.287 | 0.192 | 0.125 | 0.115 | 0.302 | 0.297 | 0.159 | 0.133 | 0.108 |
| 5% | 0.119 | 0.202 | 0.242 | 0.166 | 0.271 | 0.331 | 0.290 | 0.170 | 0.118 | 0.090 | 0.328 | 0.295 | 0.154 | 0.118 | 0.105 |
| 10% | 0.028 | 0.048 | 0.284 | 0.183 | 0.458 | 0.331 | 0.318 | 0.171 | 0.097 | 0.084 | 0.321 | 0.298 | 0.164 | 0.116 | 0.102 |
| 20% | <0.001 | 0.004 | 0.150 | 0.209 | 0.636 | 0.359 | 0.298 | 0.155 | 0.109 | 0.079 | 0.267 | 0.269 | 0.171 | 0.137 | 0.156 |
| Half-life = ∞; $\sigma_1^2 = 1, \sigma_2^2 = 3$; $\theta_0 = 1$ | | | | | | | | | | | | | | | |
| $\xi_c$=1% | <0.001 | 0.706 | <0.001 | <0.001 | 0.293 | <0.001 | 0.722 | <0.001 | <0.001 | 0.277 | <0.001 | 0.760 | <0.001 | <0.001 | 0.239 |
| 2.5% | 0.001 | 0.606 | 0.001 | <0.001 | 0.392 | 0.001 | 0.673 | 0.001 | <0.001 | 0.325 | 0.002 | 0.726 | 0.001 | 0.001 | 0.271 |
| 5% | 0.005 | 0.390 | 0.024 | 0.018 | 0.563 | 0.002 | 0.727 | 0.001 | 0.001 | 0.268 | 0.006 | 0.732 | 0.003 | 0.003 | 0.256 |
| 10% | 0.002 | 0.129 | 0.036 | 0.040 | 0.793 | 0.001 | 0.767 | 0.001 | 0.001 | 0.230 | 0.018 | 0.692 | 0.012 | 0.004 | 0.274 |
| 20% | <0.001 | 0.002 | 0.079 | 0.098 | 0.820 | 0.008 | 0.762 | 0.003 | 0.002 | 0.226 | 0.044 | 0.632 | 0.032 | 0.032 | 0.261 |
| Half-life = ∞; $\sigma_1^2 = 1, \sigma_2^2 = 5$; $\theta_0 = 1$ | | | | | | | | | | | | | | | |
| $\xi_c$=1% | <0.001 | 0.711 | <0.001 | <0.001 | 0.289 | <0.001 | 0.719 | <0.001 | <0.001 | 0.281 | <0.001 | 0.775 | <0.001 | <0.001 | 0.225 |
| 2.5% | <0.001 | 0.656 | <0.001 | <0.001 | 0.344 | <0.001 | 0.718 | <0.001 | <0.001 | 0.282 | <0.001 | 0.753 | <0.001 | <0.001 | 0.247 |
| 5% | <0.001 | 0.452 | <0.001 | <0.001 | 0.548 | <0.001 | 0.713 | <0.001 | <0.001 | 0.287 | <0.001 | 0.734 | <0.001 | <0.001 | 0.266 |
| 10% | <0.001 | 0.126 | 0.008 | 0.004 | 0.863 | <0.001 | 0.737 | <0.001 | <0.001 | 0.263 | <0.001 | 0.718 | <0.001 | <0.001 | 0.282 |

| | | | | | | | | | | | | | | | |
|---|---|---|---|---|---|---|---|---|---|---|---|---|---|---|---|
| 20% | <0.001 | 0.015 | 0.047 | 0.041 | 0.897 | <0.001 | 0.792 | <0.001 | <0.001 | 0.208 | 0.001 | 0.722 | 0.001 | 0.001 | 0.276 |
| Half-life = 0.50; $\sigma_1^2=1, \sigma_2^2=1.25$; $\theta_1=1, \theta_2=3$ | | | | | | | | | | | | | | | |
| $\xi_c$=1% | 0.011 | 0.018 | 0.020 | 0.456 | 0.495 | 0.012 | 0.019 | 0.021 | 0.476 | 0.472 | 0.014 | 0.022 | 0.014 | 0.456 | 0.494 |
| 2.5% | 0.005 | 0.014 | 0.014 | 0.347 | 0.619 | 0.012 | 0.024 | 0.021 | 0.470 | 0.472 | 0.013 | 0.029 | 0.011 | 0.431 | 0.516 |
| 5% | <0.001 | 0.001 | 0.007 | 0.221 | 0.770 | 0.010 | 0.011 | 0.015 | 0.494 | 0.470 | 0.010 | 0.014 | 0.012 | 0.448 | 0.516 |
| 10% | <0.001 | <0.001 | 0.012 | 0.151 | 0.838 | 0.024 | 0.028 | 0.023 | 0.489 | 0.436 | 0.021 | 0.035 | 0.012 | 0.340 | 0.592 |
| 20% | <0.001 | <0.001 | 0.007 | 0.153 | 0.840 | 0.036 | 0.032 | 0.017 | 0.491 | 0.425 | 0.017 | 0.032 | 0.010 | 0.320 | 0.621 |
| Half-life = 0.50; $\sigma_1^2=1, \sigma_2^2=3$; $\theta_1=1, \theta_2=3$ | | | | | | | | | | | | | | | |
| $\xi_c$=1% | <0.001 | 0.073 | <0.001 | <0.001 | 0.926 | <0.001 | 0.080 | <0.001 | 0.001 | 0.919 | <0.001 | 0.090 | <0.001 | 0.001 | 0.909 |
| 2.5% | <0.001 | 0.057 | <0.001 | 0.001 | 0.941 | <0.001 | 0.085 | <0.001 | 0.001 | 0.913 | <0.001 | 0.101 | 0.001 | 0.009 | 0.888 |
| 5% | <0.001 | 0.019 | <0.001 | 0.003 | 0.978 | <0.001 | 0.079 | <0.001 | 0.001 | 0.920 | <0.001 | 0.078 | 0.001 | 0.010 | 0.911 |
| 10% | <0.001 | 0.001 | 0.002 | 0.008 | 0.990 | <0.001 | 0.104 | <0.001 | 0.003 | 0.892 | <0.001 | 0.112 | 0.001 | 0.017 | 0.869 |
| 20% | <0.001 | <0.001 | 0.001 | 0.026 | 0.973 | <0.001 | 0.248 | <0.001 | <0.001 | 0.751 | 0.001 | 0.219 | 0.002 | 0.006 | 0.772 |
| Half-life = 0.50; $\sigma_1^2=1, \sigma_2^2=5$; $\theta_1=1, \theta_2=3$ | | | | | | | | | | | | | | | |
| $\xi_c$=1% | <0.001 | 0.105 | <0.001 | <0.001 | 0.895 | <0.001 | 0.118 | <0.001 | <0.001 | 0.882 | <0.001 | 0.224 | <0.001 | <0.001 | 0.776 |
| 2.5% | <0.001 | 0.063 | <0.001 | <0.001 | 0.937 | <0.001 | 0.099 | <0.001 | <0.001 | 0.901 | <0.001 | 0.126 | <0.001 | <0.001 | 0.874 |
| 5% | <0.001 | 0.033 | <0.001 | <0.001 | 0.967 | <0.001 | 0.156 | <0.001 | <0.001 | 0.844 | <0.001 | 0.185 | <0.001 | <0.001 | 0.815 |
| 10% | <0.001 | 0.002 | <0.001 | <0.001 | 0.998 | <0.001 | 0.221 | <0.001 | <0.001 | 0.779 | <0.001 | 0.192 | <0.001 | <0.001 | 0.808 |
| 20% | <0.001 | <0.001 | 0.001 | 0.002 | 0.997 | <0.001 | 0.283 | <0.001 | <0.001 | 0.717 | <0.001 | 0.218 | <0.001 | <0.001 | 0.782 |
| Half-life = 0.50; $\sigma_1^2=2, \sigma_2^2=2$; $\theta_1=1, \theta_2=1.25$ | | | | | | | | | | | | | | | |
| $\xi_c$=1% | 0.038 | 0.043 | 0.415 | 0.302 | 0.202 | 0.039 | 0.043 | 0.416 | 0.300 | 0.202 | 0.172 | 0.136 | 0.193 | 0.320 | 0.180 |
| 2.5% | 0.018 | 0.050 | 0.433 | 0.305 | 0.194 | 0.021 | 0.050 | 0.434 | 0.302 | 0.193 | 0.154 | 0.138 | 0.263 | 0.307 | 0.139 |
| 5% | 0.028 | 0.055 | 0.390 | 0.306 | 0.221 | 0.034 | 0.062 | 0.397 | 0.300 | 0.207 | 0.142 | 0.133 | 0.241 | 0.324 | 0.160 |
| 10% | 0.014 | 0.016 | 0.395 | 0.345 | 0.230 | 0.034 | 0.033 | 0.392 | 0.332 | 0.208 | 0.143 | 0.128 | 0.203 | 0.341 | 0.184 |
| 20% | 0.012 | 0.017 | 0.382 | 0.324 | 0.265 | 0.067 | 0.089 | 0.389 | 0.269 | 0.186 | 0.178 | 0.163 | 0.233 | 0.260 | 0.166 |
| Half-life = 0.50; $\sigma_1^2=2, \sigma_2^2=2$; $\theta_1=1, \theta_2=3$ | | | | | | | | | | | | | | | |
| $\xi_c$=1% | 0.001 | 0.001 | 0.006 | 0.634 | 0.358 | 0.001 | 0.001 | 0.006 | 0.638 | 0.354 | 0.006 | 0.007 | 0.004 | 0.626 | 0.357 |
| 2.5% | <0.001 | 0.001 | 0.010 | 0.566 | 0.423 | <0.001 | 0.001 | 0.010 | 0.568 | 0.420 | 0.002 | 0.003 | 0.001 | 0.611 | 0.383 |
| 5% | <0.001 | 0.001 | 0.002 | 0.611 | 0.387 | 0.001 | 0.001 | 0.002 | 0.610 | 0.385 | 0.015 | 0.016 | 0.009 | 0.612 | 0.348 |

| | | | | | | | | | | | | | | | |
|---|---|---|---|---|---|---|---|---|---|---|---|---|---|---|---|
| 10% | <0.001 | <0.001 | 0.002 | 0.620 | 0.379 | 0.001 | <0.001 | 0.002 | 0.627 | 0.369 | 0.007 | 0.004 | 0.004 | 0.659 | 0.326 |
| 20% | 0.002 | 0.001 | 0.004 | 0.520 | 0.473 | 0.005 | 0.003 | 0.006 | 0.616 | 0.370 | 0.025 | 0.016 | 0.011 | 0.537 | 0.411 |
| Half-life = 0.50; $\sigma_1^2 = 2, \sigma_2^2 = 2$; $\theta_1 = 1, \theta_2 = 5$ | | | | | | | | | | | | | | | |
| $\xi_c$=1% | <0.001 | <0.001 | <0.001 | 0.633 | 0.367 | <0.001 | <0.001 | <0.001 | 0.639 | 0.361 | <0.001 | <0.001 | <0.001 | 0.698 | 0.302 |
| 2.5% | <0.001 | <0.001 | <0.001 | 0.625 | 0.375 | <0.001 | <0.001 | <0.001 | 0.625 | 0.375 | <0.001 | <0.001 | <0.001 | 0.653 | 0.347 |
| 5% | <0.001 | <0.001 | <0.001 | 0.607 | 0.393 | <0.001 | <0.001 | <0.001 | 0.641 | 0.359 | <0.001 | <0.001 | <0.001 | 0.671 | 0.329 |
| 10% | <0.001 | <0.001 | <0.001 | 0.561 | 0.439 | <0.001 | <0.001 | <0.001 | 0.607 | 0.393 | <0.001 | <0.001 | <0.001 | 0.606 | 0.394 |
| 20% | <0.001 | <0.001 | <0.001 | 0.315 | 0.685 | <0.001 | <0.001 | <0.001 | 0.628 | 0.372 | <0.001 | <0.001 | <0.001 | 0.389 | 0.611 |

BM1 = Single rate BM; BMS = Multiple rate BM; OU1 = single optimum OU; OUM = multiple optima OU; OUMV = multiple optima and rate OU.

**Table S2.** Summary of model support, parameter estimates, tip fog, and various other summary statistics obtained from a set of empirical data sets.

| Dataset | Model | Fog | Program | ΔAICc | Tip fog as percentage of tip mean | Percent of variance from tip fog | $\sigma^2$ | $\xi_c$ | Other *geiger* parameter | Number of taxa | Original Tree Height |
|---|---|---|---|---|---|---|---|---|---|---|---|
| **AlencarEtAl_log_precip** | **kappa** | **none** | **geiger** | **0.00** | **0%** | **0%** | **0.15** | **0.00** | **0.08** | **663** | **83.49** |
| AlencarEtAl_log_precip | kappa | estimate | geiger | 0.88 | 8% | 14% | 0.14 | 0.25 | 0.00 | 663 | 83.49 |
| AlencarEtAl_log_precip | EB | estimate | geiger | 47.95 | 10% | 26% | 0.04 | 0.47 | -0.02 | 663 | 83.49 |
| AlencarEtAl_log_precip | BM1 | estimate | ouwie | 48.92 | 7% | | 0.01 | 0.20 | | 663 | 83.49 |
| AlencarEtAl_log_precip | delta | estimate | geiger | 48.92 | 10% | 26% | 0.02 | 0.46 | 0.53 | 663 | 83.49 |
| AlencarEtAl_log_precip | OU1 | estimate | ouwie | 50.94 | 7% | | 0.01 | 0.20 | | 663 | 83.49 |
| AlencarEtAl_log_precip | OU | none | geiger | 249.46 | 0% | 0% | 0.09 | 0.00 | 0.04 | 663 | 83.49 |
| AlencarEtAl_log_precip | OU | estimate | geiger | 249.46 | 0% | 0% | 0.09 | 0.00 | 0.04 | 663 | 83.49 |
| AlencarEtAl_log_precip | OU1 | none | ouwie | 249.46 | 0% | | 0.09 | 0.00 | | 663 | 83.49 |
| AlencarEtAl_log_precip | delta | none | geiger | 325.91 | 0% | 0% | 0.02 | 0.00 | 3.00 | 663 | 83.49 |
| AlencarEtAl_log_precip | BM | none | geiger | 416.81 | 0% | 0% | 0.06 | 0.00 | | 663 | 83.49 |
| AlencarEtAl_log_precip | BM1 | none | ouwie | 416.81 | 0% | | 0.06 | 0.00 | | 663 | 83.49 |
| AlencarEtAl_log_precip | BM | estimate | geiger | 418.83 | 0% | 0% | 0.06 | 0.00 | | 663 | 83.49 |
| AlencarEtAl_log_precip | EB | none | geiger | 418.83 | 0% | 0% | 0.06 | 0.00 | 0.00 | 663 | 83.49 |
| AlencarEtAl_log_precip | white | only | geiger | 765.24 | 18% | 100% | 0.00 | 1.44 | | 663 | 83.49 |
| **AlencarEtAl_log_svl** | **EB** | **estimate** | **geiger** | **0.00** | **8%** | **37%** | **0.01** | **0.11** | **-0.02** | **596** | **83.49** |
| AlencarEtAl_log_svl | delta | estimate | geiger | 1.89 | 8% | 36% | 0.00 | 0.10 | 0.46 | 596 | 83.49 |
| AlencarEtAl_log_svl | BM | estimate | geiger | 3.42 | 7% | 45% | 0.00 | 0.10 | | 596 | 83.49 |
| AlencarEtAl_log_svl | BM1 | estimate | ouwie | 3.42 | 2% | | 0.00 | 0.01 | | 596 | 83.49 |
| AlencarEtAl_log_svl | OU1 | estimate | ouwie | 5.45 | 2% | | 0.00 | 0.01 | | 596 | 83.49 |
| AlencarEtAl_log_svl | kappa | none | geiger | 21.42 | 0% | 0% | 0.01 | 0.00 | 0.47 | 596 | 83.49 |

| | | | | | | | | | | | |
|---|---|---|---|---|---|---|---|---|---|---|---|
| AlencarEtAl_log_svl | kappa | estimate | geiger | 23.45 | 0% | 0% | 0.01 | 0.00 | 0.47 | 596 | 83.49 |
| AlencarEtAl_log_svl | OU | none | geiger | 109.13 | 0% | 0% | 0.00 | 0.00 | 0.01 | 596 | 83.49 |
| AlencarEtAl_log_svl | OU | estimate | geiger | 109.13 | 0% | 0% | 0.00 | 0.00 | 0.01 | 596 | 83.49 |
| AlencarEtAl_log_svl | OU1 | none | ouwie | 109.13 | 0% | | 0.00 | 0.00 | | 596 | 83.49 |
| AlencarEtAl_log_svl | delta | none | geiger | 122.45 | 0% | 0% | 0.00 | 0.00 | 2.27 | 596 | 83.49 |
| AlencarEtAl_log_svl | BM1 | none | ouwie | 141.63 | 0% | | 0.00 | 0.00 | | 596 | 83.49 |
| AlencarEtAl_log_svl | BM | none | geiger | 141.63 | 0% | 0% | 0.00 | 0.00 | | 596 | 83.49 |
| AlencarEtAl_log_svl | EB | none | geiger | 143.66 | 0% | 0% | 0.00 | 0.00 | 0.00 | 596 | 83.49 |
| AlencarEtAl_log_svl | white | only | geiger | 908.75 | 10% | 100% | 0.00 | 0.19 | | 596 | 83.49 |
| **AlencarEtAl_log_tempK** | **kappa** | **estimate** | **geiger** | **0.00** | **1%** | **96%** | **0.00** | **0.01** | **0.00** | **663** | **83.49** |
| AlencarEtAl_log_tempK | BM | estimate | geiger | 17.68 | 2% | 97% | 0.00 | 0.01 | | 663 | 83.49 |
| AlencarEtAl_log_tempK | BM1 | estimate | ouwie | 17.68 | 0% | | 0.00 | 0.00 | | 663 | 83.49 |
| AlencarEtAl_log_tempK | EB | estimate | geiger | 19.70 | 2% | 97% | 0.00 | 0.01 | 0.00 | 663 | 83.49 |
| AlencarEtAl_log_tempK | delta | estimate | geiger | 19.71 | 2% | 97% | 0.00 | 0.01 | 0.98 | 663 | 83.49 |
| AlencarEtAl_log_tempK | OU1 | estimate | ouwie | 19.71 | 0% | | 0.00 | 0.00 | | 663 | 83.49 |
| AlencarEtAl_log_tempK | kappa | none | geiger | 26.44 | 0% | 0% | 0.00 | 0.00 | 0.09 | 663 | 83.49 |
| AlencarEtAl_log_tempK | OU | none | geiger | 164.52 | 0% | 0% | 0.00 | 0.00 | 0.04 | 663 | 83.49 |
| AlencarEtAl_log_tempK | OU | estimate | geiger | 164.52 | 0% | 0% | 0.00 | 0.00 | 0.04 | 663 | 83.49 |
| AlencarEtAl_log_tempK | OU1 | none | ouwie | 164.52 | 0% | | 0.00 | 0.00 | | 663 | 83.49 |
| AlencarEtAl_log_tempK | delta | none | geiger | 232.62 | 0% | 0% | 0.00 | 0.00 | 3.00 | 663 | 83.49 |
| AlencarEtAl_log_tempK | BM | none | geiger | 322.91 | 0% | 0% | 0.00 | 0.00 | | 663 | 83.49 |
| AlencarEtAl_log_tempK | BM1 | none | ouwie | 322.91 | 0% | | 0.00 | 0.00 | | 663 | 83.49 |
| AlencarEtAl_log_tempK | EB | none | geiger | 324.94 | 0% | 0% | 0.00 | 0.00 | 0.00 | 663 | 83.49 |
| AlencarEtAl_log_tempK | white | only | geiger | 845.82 | 0% | 100% | 0.00 | 0.00 | | 663 | 83.49 |
| **AlencarEtAl_log_topo_complexity** | **kappa** | **estimate** | **geiger** | **0.00** | **47%** | **44%** | **0.06** | **0.74** | **0.18** | **662** | **83.49** |
| AlencarEtAl_log_topo_complexity | BM | estimate | geiger | 14.08 | 48% | 51% | 0.01 | 0.78 | | 662 | 83.49 |
| AlencarEtAl_log_topo_complexity | BM1 | estimate | ouwie | 14.08 | 42% | | 0.01 | 0.61 | | 662 | 83.49 |
| AlencarEtAl_log_topo_complexity | OU1 | estimate | ouwie | 14.83 | 42% | | 0.01 | 0.58 | | 662 | 83.49 |
| AlencarEtAl_log_topo_complexity | EB | estimate | geiger | 16.10 | 48% | 51% | 0.01 | 0.78 | 0.00 | 662 | 83.49 |

| | | | | | | | | | | | |
|---|---|---|---|---|---|---|---|---|---|---|---|
| AlencarEtAl_log_topo_complexity | kappa | none | geiger | 89.63 | 0% | 0% | 0.40 | 0.00 | 0.00 | 662 | 83.49 |
| AlencarEtAl_log_topo_complexity | OU | none | geiger | 131.18 | 0% | 0% | 0.40 | 0.00 | 0.17 | 662 | 83.49 |
| AlencarEtAl_log_topo_complexity | OU | estimate | geiger | 131.18 | 0% | 0% | 0.40 | 0.00 | 0.17 | 662 | 83.49 |
| AlencarEtAl_log_topo_complexity | OU1 | none | ouwie | 131.18 | 0% | | 0.40 | 0.00 | | 662 | 83.49 |
| AlencarEtAl_log_topo_complexity | white | only | geiger | 169.32 | 59% | 100% | 0.00 | 1.16 | | 662 | 83.49 |
| AlencarEtAl_log_topo_complexity | delta | estimate | geiger | 173.37 | 57% | 100% | 0.00 | 1.07 | 0.00 | 662 | 83.49 |
| AlencarEtAl_log_topo_complexity | delta | none | geiger | 382.39 | 0% | 0% | 0.05 | 0.00 | 3.00 | 662 | 83.49 |
| AlencarEtAl_log_topo_complexity | BM | none | geiger | 502.36 | 0% | 0% | 0.13 | 0.00 | | 662 | 83.49 |
| AlencarEtAl_log_topo_complexity | BM1 | none | ouwie | 502.36 | 0% | | 0.13 | 0.00 | | 662 | 83.49 |
| AlencarEtAl_log_topo_complexity | EB | none | geiger | 504.38 | 0% | 0% | 0.13 | 0.00 | 0.00 | 662 | 83.49 |
| **caniformia** | **kappa** | **none** | **geiger** | **0.00** | **0%** | **0%** | **0.16** | **0.00** | **0.58** | **135** | **48.90** |
| caniformia | kappa | estimate | geiger | 2.12 | 0% | 0% | 0.16 | 0.00 | 0.58 | 135 | 48.90 |
| caniformia | BM | estimate | geiger | 5.94 | 21% | 6% | 0.10 | 0.23 | | 135 | 48.90 |
| caniformia | BM1 | estimate | ouwie | 5.94 | 10% | | 0.10 | 0.05 | | 135 | 48.90 |
| caniformia | delta | estimate | geiger | 8.07 | 21% | 6% | 0.10 | 0.23 | 0.98 | 135 | 48.90 |
| caniformia | OU1 | estimate | ouwie | 8.07 | 10% | | 0.10 | 0.05 | | 135 | 48.90 |
| caniformia | OU | estimate | geiger | 8.07 | 21% | 6% | 0.10 | 0.23 | 0.00 | 135 | 48.90 |
| caniformia | BM1 | none | ouwie | 10.49 | 0% | | 0.12 | 0.00 | | 135 | 48.90 |
| caniformia | BM | none | geiger | 10.49 | 0% | 0% | 0.12 | 0.00 | | 135 | 48.90 |
| caniformia | OU | none | geiger | 11.93 | 0% | 0% | 0.13 | 0.00 | 0.01 | 135 | 48.90 |
| caniformia | OU1 | none | ouwie | 11.93 | 0% | | 0.13 | 0.00 | | 135 | 48.90 |
| caniformia | delta | none | geiger | 12.11 | 0% | 0% | 0.09 | 0.00 | 1.40 | 135 | 48.90 |
| caniformia | EB | none | geiger | 12.59 | 0% | 0% | 0.12 | 0.00 | 0.00 | 135 | 48.90 |
| caniformia | EB | estimate | geiger | 14.71 | 0% | 0% | 0.12 | 0.00 | 0.00 | 135 | 48.90 |
| caniformia | white | only | geiger | 266.17 | 95% | 100% | 0.00 | 4.73 | | 135 | 48.90 |
| **carnivores** | **BM** | **none** | **geiger** | **0.00** | **0%** | **0%** | **0.07** | **0.00** | | **16** | **59.20** |
| carnivores | BM1 | none | ouwie | 0.00 | 0% | | 0.07 | 0.00 | | 16 | 59.20 |
| carnivores | kappa | none | geiger | 2.18 | 0% | 0% | 0.39 | 0.00 | 0.44 | 16 | 59.20 |
| carnivores | delta | none | geiger | 3.00 | 0% | 0% | 0.07 | 0.00 | 1.25 | 16 | 59.20 |

| | | | | | | | | | | | |
|---|---|---|---|---|---|---|---|---|---|---|---|
| carnivores | EB | none | geiger | 3.07 | 0% | 0% | 0.08 | 0.00 | 0.00 | 16 | 59.20 |
| carnivores | OU1 | none | ouwie | 3.08 | 0% | | 0.07 | 0.00 | | 16 | 59.20 |
| carnivores | BM | estimate | geiger | 3.08 | 0% | 0% | 0.07 | 0.00 | | 16 | 59.20 |
| carnivores | OU | none | geiger | 3.08 | 0% | | 0.07 | 0.00 | 0.00 | 16 | 59.20 |
| carnivores | BM1 | estimate | ouwie | 3.08 | 0% | | 0.07 | 0.00 | | 16 | 59.20 |
| carnivores | white | only | geiger | 4.89 | 98% | 100% | 0.00 | 4.51 | | 16 | 59.20 |
| carnivores | kappa | estimate | geiger | 5.81 | 0% | 0% | 0.39 | 0.00 | 0.44 | 16 | 59.20 |
| carnivores | delta | estimate | geiger | 6.64 | 0% | 0% | 0.07 | 0.00 | 1.25 | 16 | 59.20 |
| carnivores | EB | estimate | geiger | 6.71 | 0% | 0% | 0.08 | 0.00 | 0.00 | 16 | 59.20 |
| carnivores | OU1 | estimate | ouwie | 6.71 | 0% | | 0.07 | 0.00 | | 16 | 59.20 |
| carnivores | OU | estimate | geiger | 6.71 | 0% | 0% | 0.07 | 0.00 | 0.00 | 16 | 59.20 |
| **geospiza** | **white** | **only** | **geiger** | **0.00** | **3%** | **100%** | **0.00** | **0.01** | | **13** | **0.58** |
| geospiza | BM | none | geiger | 3.12 | 0% | 0% | 0.07 | 0.00 | | 13 | 0.58 |
| geospiza | BM1 | none | ouwie | 3.12 | 0% | | 0.07 | 0.00 | | 13 | 0.58 |
| geospiza | OU1 | none | ouwie | 3.39 | 0% | | 0.42 | 0.00 | | 13 | 0.58 |
| geospiza | BM | estimate | geiger | 3.47 | 8% | 100% | 0.00 | 0.11 | | 13 | 0.58 |
| geospiza | BM1 | estimate | ouwie | 3.62 | 2% | | 0.02 | 0.01 | | 13 | 0.58 |
| geospiza | OU | none | geiger | 4.20 | 0% | 0% | 0.11 | 0.00 | 2.72 | 13 | 0.58 |
| geospiza | delta | none | geiger | 4.35 | 0% | 0% | 0.04 | 0.00 | 3.00 | 13 | 0.58 |
| geospiza | kappa | none | geiger | 5.30 | 0% | 0% | 0.03 | 0.00 | 0.61 | 13 | 0.58 |
| geospiza | EB | none | geiger | 6.59 | 0% | 0% | 0.07 | 0.00 | 0.00 | 13 | 0.58 |
| geospiza | EB | estimate | geiger | 7.80 | 8% | 100% | 0.00 | 0.11 | -8.70 | 13 | 0.58 |
| geospiza | kappa | estimate | geiger | 7.80 | 8% | 100% | 0.00 | 0.11 | 0.00 | 13 | 0.58 |
| geospiza | OU | estimate | geiger | 7.80 | 8% | 100% | 0.00 | 0.11 | 0.00 | 13 | 0.58 |
| geospiza | delta | estimate | geiger | 7.80 | 8% | 100% | 0.00 | 0.11 | 0.00 | 13 | 0.58 |
| geospiza | OU1 | estimate | ouwie | 7.80 | 3% | | 0.00 | 0.01 | | 13 | 0.58 |
| **primates** | **EB** | **estimate** | **geiger** | **0.00** | **4%** | **4%** | **0.12** | **0.10** | **-0.03** | **233** | **65.09** |
| primates | BM1 | estimate | ouwie | 1.84 | 1% | | 0.03 | 0.01 | | 233 | 65.09 |
| primates | OU | estimate | geiger | 3.91 | 4% | 100% | 0.03 | 0.08 | 0.00 | 233 | 65.09 |

| | | | | | | | | | | | |
|---|---|---|---|---|---|---|---|---|---|---|---|
| primates | OU1 | estimate | ouwie | 3.91 | 1% | | 0.03 | 0.01 | | 233 | 65.09 |
| primates | kappa | none | geiger | 5.08 | 0% | 0% | 0.04 | 0.00 | 0.87 | 233 | 65.09 |
| primates | BM1 | none | ouwie | 5.72 | 0% | | 0.03 | 0.00 | | 233 | 65.09 |
| primates | BM | none | geiger | 5.72 | 0% | 0% | 0.03 | 0.00 | | 233 | 65.09 |
| primates | EB | none | geiger | 6.87 | 0% | 0% | 0.06 | 0.00 | -0.01 | 233 | 65.09 |
| primates | delta | none | geiger | 6.98 | 0% | 0% | 0.05 | 0.00 | 0.67 | 233 | 65.09 |
| primates | kappa | estimate | geiger | 7.15 | 0% | 0% | 0.04 | 0.00 | 0.87 | 233 | 65.09 |
| primates | OU1 | none | ouwie | 7.77 | 0% | | 0.03 | 0.00 | | 233 | 65.09 |
| primates | OU | none | geiger | 7.77 | 0% | | 0.03 | 0.00 | 0.00 | 233 | 65.09 |
| primates | BM | estimate | geiger | 7.77 | 0% | 0% | 0.03 | 0.00 | | 233 | 65.09 |
| primates | delta | estimate | geiger | 9.05 | 0% | 0% | 0.05 | 0.00 | 0.67 | 233 | 65.09 |
| primates | white | only | geiger | 550.78 | 19% | 100% | 0.00 | 2.08 | | 233 | 65.09 |

**Table S3.** Summary of model support for simulated scenarios of discrete-trait evolution that contained increasing levels of tip fog, assessed by calculating the average Akaike weight ($w$) for all models assessed the fit.

| | Default | | $+\ \xi_d$ | | Pooled model set | | | |
|---|---|---|---|---|---|---|---|---|
| Model | $ER_w$ | $ARD_w$ | $ER_w$ | $ARD_w$ | $ER_w$ | $ARD_w$ | $ER_w + \xi_d$ | $ARD_w + \xi_d$ |
| Clade Age = 5 Myr | | | | | | | | |
| $\xi_d$ = 1% | 0.532 | 0.468 | 0.605 | 0.395 | 0.221 | 0.257 | 0.243 | 0.188 |
| 5% | 0.014 | 0.986 | 0.577 | 0.423 | <0.001 | 0.041 | 0.446 | 0.302 |
| 10% | <0.001 | 0.999 | 0.576 | 0.424 | <0.001 | 0.091 | 0.437 | 0.318 |
| 15% | <0.001 | 0.999 | 0.576 | 0.424 | <0.001 | 0.133 | 0.417 | 0.328 |
| 20% | <0.001 | 0.999 | 0.576 | 0.424 | <0.001 | 0.143 | 0.426 | 0.304 |
| Clade age = 10 Myr | | | | | | | | |
| $\xi_d$ = 1% | 0.604 | 0.396 | 0.638 | 0.362 | 0.293 | 0.231 | 0.216 | 0.165 |
| 5% | 0.100 | 0.900 | 0.653 | 0.347 | <0.001 | 0.008 | 0.608 | 0.317 |
| 10% | 0.003 | 0.997 | 0.635 | 0.365 | <0.001 | <0.001 | 0.625 | 0.333 |
| 15% | <0.001 | 0.999 | 0.576 | 0.424 | <0.001 | 0.006 | 0.525 | 0.323 |
| 20% | <0.001 | 0.999 | 0.593 | 0.407 | <0.001 | 0.012 | 0.530 | 0.322 |
| Clade age = 15 Myr | | | | | | | | |
| $\xi_d$ = 1% | 0.616 | 0.384 | 0.658 | 0.342 | 0.319 | 0.206 | 0.225 | 0.158 |
| 5% | 0.465 | 0.535 | 0.645 | 0.355 | 0.001 | 0.006 | 0.599 | 0.304 |
| 10% | 0.064 | 0.936 | 0.643 | 0.357 | <0.001 | <0.001 | 0.635 | 0.325 |
| 15% | <0.001 | 0.999 | 0.659 | 0.341 | <0.001 | <0.001 | 0.658 | 0.317 |
| 20% | <0.001 | 0.999 | 0.617 | 0.383 | <0.001 | 0.001 | 0.606 | 0.304 |
| Clade age = 20 Myr | | | | | | | | |
| $\xi_d$ = 1% | 0.663 | 0.337 | 0.685 | 0.315 | 0.332 | 0.198 | 0.246 | 0.146 |
| 5% | 0.480 | 0.520 | 0.680 | 0.320 | 0.002 | 0.009 | 0.604 | 0.294 |
| 10% | 0.304 | 0.696 | 0.665 | 0.335 | <0.001 | <0.001 | 0.663 | 0.327 |
| 15% | <0.001 | 0.999 | 0.627 | 0.373 | <0.001 | <0.001 | 0.622 | 0.341 |
| 20% | <0.001 | 0.999 | 0.642 | 0.358 | <0.001 | 0.001 | 0.626 | 0.309 |

ER = Equal rates; ARD = All rates different

**Table S4.** Same simulation scenario as presented in Table S2, but in this case we added an HMM into the model set and recalculated the average Akaike weight ($w$) to assess fit.

| | Default | | | $+ \xi_d$ | | |
|---|---|---|---|---|---|---|
| Model | $ER_w$ | $ARD_w$ | $HMM_w$ | $ER_w$ | $ARD_w$ | $HMM_w$ |
| Clade age = 5 Myr | | | | | | |
| $\xi_d$ = 1% | 0.465 | 0.437 | 0.071 | 0.559 | 0.365 | 0.076 |
| 5% | 0.005 | 0.839 | 0.051 | 0.535 | 0.393 | 0.076 |
| 10% | <0.001 | 0.999 | 0.001 | 0.534 | 0.393 | 0.072 |
| 15% | <0.001 | 0.999 | <0.001 | 0.534 | 0.393 | 0.072 |
| 20% | <0.001 | 0.999 | <0.001 | 0.534 | 0.394 | 0.072 |
| Clade age = 10 Myr | | | | | | |
| $\xi_d$ = 1% | 0.533 | 0.353 | 0.082 | 0.582 | 0.329 | 0.083 |
| 5% | 0.033 | 0.401 | 0.162 | 0.596 | 0.318 | 0.083 |
| 10% | <0.001 | 0.303 | 0.687 | 0.581 | 0.331 | 0.082 |
| 15% | <0.001 | 0.962 | 0.038 | 0.534 | 0.391 | 0.076 |
| 20% | <0.001 | 0.998 | 0.002 | 0.534 | 0.376 | 0.080 |
| Clade age = 15 Myr | | | | | | |
| $\xi_d$ = 1% | 0.532 | 0.309 | 0.088 | 0.600 | 0.305 | 0.086 |
| 5% | 0.094 | 0.249 | 0.245 | 0.575 | 0.322 | 0.082 |
| 10% | <0.001 | 0.055 | 0.925 | 0.587 | 0.328 | 0.083 |
| 15% | <0.001 | 0.068 | 0.927 | 0.597 | 0.304 | 0.087 |
| 20% | <0.001 | 0.677 | 0.323 | 0.561 | 0.344 | 0.083 |
| Clade age = 20 Myr | | | | | | |
| $\xi_d$ = 1% | 0.595 | 0.308 | 0.088 | 0.614 | 0.287 | 0.088 |
| 5% | 0.220 | 0.303 | 0.215 | 0.610 | 0.293 | 0.087 |
| 10% | 0.001 | 0.018 | 0.966 | 0.603 | 0.303 | 0.085 |
| 15% | <0.001 | 0.060 | 0.901 | 0.567 | 0.331 | 0.085 |
| 20% | <0.001 | 0.206 | 0.794 | 0.563 | 0.322 | 0.085 |

ER = Equal rates; ARD = All rates different; HMM = hidden rates model

**Table S5.** Summary of model support for simulated scenarios of character-dependent diversification that contained increasing levels of tip fog, assessed by calculating the average Akaike weight ($w$) for all models assessed the fit.

| Model | Type | $\xi_d$ | | | | |
|---|---|---|---|---|---|---|
| | | 1% | 5% | 10% | 15% | 20% |
| $\lambda_0 = \lambda_1, \mu_0 = \mu_1, q_{01} = q_{10}$ | CID | 0.124 | 0.112 | 0.102 | 0.109 | 0.126 |
| $\lambda_0 = \lambda_1, \mu_0 = \mu_1, q_{01} \neq q_{10}$ | CID | 0.093 | 0.108 | 0.133 | 0.203 | 0.251 |
| $\lambda_0 \neq \lambda_1, \mu_0 \neq \mu_1, q_{01} = q_{10}$ | CD | 0.445 | 0.382 | 0.299 | 0.202 | 0.145 |
| $\lambda_0 \neq \lambda_1, \mu_0 \neq \mu_1, q_{01} \neq q_{10}$ | CD | 0.257 | 0.302 | 0.368 | 0.356 | 0.294 |
| $\lambda_{0A} = \lambda_{1A} \neq \lambda_{0B} = \lambda_{1B}, \mu_{0A} = \lambda\mu_{1A} \neq \mu_{0B} = \lambda\mu_{1B}, q_{0A1A} = q_{0B1B}$ | CID | 0.056 | 0.058 | 0.060 | 0.052 | 0.056 |
| $\lambda_{0A} = \lambda_{1A} \neq \lambda_{0B} = \lambda_{1B}, \mu_{0A} = \lambda\mu_{1A} \neq \mu_{0B} = \lambda\mu_{1B}, q_{0A1A} \neq q_{0B1B}$ | CID | 0.025 | 0.038 | 0.038 | 0.078 | 0.128 |
| $\lambda_0 = \lambda_1, \mu_0 = \mu_1, q_{01} = q_{10}, +\xi_d$ | CID | 0.131 | 0.136 | 0.148 | 0.157 | 0.165 |
| $\lambda_0 = \lambda_1, \mu_0 = \mu_1, q_{01} \neq q_{10}, +\xi_d$ | CID | 0.084 | 0.089 | 0.094 | 0.096 | 0.092 |
| $\lambda_0 \neq \lambda_1, \mu_0 \neq \mu_1, q_{01} = q_{10}, +\xi_d$ | CD | 0.444 | 0.429 | 0.398 | 0.379 | 0.378 |
| $\lambda_0 \neq \lambda_1, \mu_0 \neq \mu_1, q_{01} \neq q_{10}, +\xi_d$ | CD | 0.251 | 0.254 | 0.261 | 0.261 | 0.257 |
| $\lambda_{0A} = \lambda_{1A} \neq \lambda_{0B} = \lambda_{1B}, \mu_{0A} = \lambda\mu_{1A} \neq \mu_{0B} = \lambda\mu_{1B}, q_{0A1A} = q_{0B1B}, +\xi_d$ | CID | 0.063 | 0.065 | 0.075 | 0.073 | 0.087 |
| $\lambda_{0A} = \lambda_{1A} \neq \lambda_{0B} = \lambda_{1B}, \mu_{0A} = \lambda\mu_{1A} \neq \mu_{0B} = \lambda\mu_{1B}, q_{0A1A} \neq q_{0B1B}, +\xi_d$ | CID | 0.026 | 0.028 | 0.024 | 0.033 | 0.022 |

CID = character-independent model; CD = character-dependent model

### C. Empirical tip fog estimates

In an earlier draft of the manuscript, we followed Silvestro et al. (2015) and used tip fog of up to 50% in our simulations. A colleague pointed out that this amount of fog represented a substantial amount of uncertainty, especially as continuous traits are often analyzed on a log scale. For example, for a mammal with body mass of 50 kg (say, a human), its log mass is 3.91 log kg, so 50% fog would mean its log mass could be from 1.96 log kg to 5.87 log kg, or from 7 to 352 kg. That is quite a lot of variation, equivalent to not being sure if a mammal has the mass of a howler monkey or polar bear.

Tip fog encompasses more than just sample variance, so rather than looking at empirical distributions of the latter, we decided to estimate tip fog directly from empirical data sets to better inform our simulations. We used datasets included in *geiger* (Pennell et al., 2014) as well as data set from a recent paper (Alencar et al. 2024). The *geiger* datasets were 'canifornmia' (Slater et al. 2012), 'primates' (Redding et al. 2010; Vos & Mooers 2006), 'carnivores' (Eizirik et al. 2010; Wozencraft, 2005; Jones et al. 2009), and 'geospiza' (Dolph Shluter data in the *geiger* package). The first two datasets have log mass. For carnivores, we used the mean column for log mass; for the geospiza dataset, "wingL" which is log wing length. For the Alencar et al. (2024) data, following their analysis we converted temperature to Kelvin and then used log transformed values of Kelvin temperature, precipitation, terrain ruggedness, and female snout vent length.

Analyses were run by fitting *geiger* models BM (Brownian motion), OU (Ornstein-Uhlenbeck), EB (early burst), kappa, delta, and white. Other than BM and white, these are single parameter transformations of the tree and then data are fit to the transformed tree essentially with a Brownian motion model. BM just uses the standard Brownian motion model. White is a white noise model, essentially converting the tree to a star phylogeny of height 1. For all but the white model (which is identical in fit to a pure tip fog only model), we also used the standard zero tip fog or allowed *geiger* to estimate it (SE=NA setting). We also fit the univariate BM1 and OU1 models in *OUwie* with and without estimated tip fog. The *OUwie* models should be equivalent to the equivalent *geiger* models; this is a way of checking one with the other. For each *geiger* model, we used the fit parameters to transform the tree, convert this to a variance-covariance matrix, and used Eq. (1) of O'Meara et al. (2006) with the model's estimated $\sigma^2$ to get expected variance at the tips from the evolutionary process. The variance at the tips from tip fog is just SE from *geiger*. We then divided the tip fog variance by the sum of both variances to calculate the percentage of species variance the model assigned to the tip fog process. We also compared the square root of the sigma.sq.me from *geiger* with the raw trait mean to compute the tip fog in units of percentage of trait mean.

Summary of the results are provided in Table S1. Taking the best (lowest AICc) model from each dataset, three datasets preferred models with no tip fog (Alencar et al. (2024) log precipitation, caniformia and carnivore body size); four estimated tip fog (Alencar et al. (2024) log snout vent length, log temperature in Kelvin, log topographic complexity; primate body size); and one chose the white model which is all tip fog (geospiza wing length). For the ones estimating tip fog, its estimate as a percentage of the mean ranged from just 1% [Alencar et al. (2024) log temperature in Kelvin] to 47% [Alencar et al. (2024) log topographic complexity], a wider range than the 0 to 20% used in our simulations. Interestingly, the percentage of all variance coming from tip fog was even higher, ranging from 4% (primate body mass) to 96% [Alencar et al. (2024) log temperature in Kelvin]. Thus, there is a wide range of empirical estimates of tip fog values, but some of the estimates can be high.

For corresponding models, *geiger* and *OUwie* returned the same AICc and parameter estimates in nearly all cases; the exceptions are some of the ones with estimated tip fog, especially larger datasets, where the *geiger* results were sometimes tens to hundreds worse in AICc. This suggests a potential issue with optimization in *geiger*, at least with default settings.

One interesting thing is that the *geiger* carnivores dataset has both mean and variance. Across the terminals, the median value for square root of the variance as a percentage of the mean (which is mean of log body mass) was 55.4% (excluding taxa with one measured individual) but went up to 288.6% (showing the magnitude of some of the uncertainty at the tips). The best fitting model, as well as the next best four models (ΔAICc of 0 to 3.08) were models that fix tip fog at zero; even among the models that could estimate tip fog, they all estimated it as zero (the exception is the “white” model, where it is forced to treat all variation as tip fog). This is a small dataset of just 16 mammal families as terminal taxa, but it shows an example of where there undoubtedly is uncertainty in the “evolutionarily true” value of log body mass for a family and yet model selection and parameter estimation did not show evidence of tip fog.

**D. References**


Alencar, L.R.V., Schwery, O., Gade, M. R., Domínguez-Guerrero, S. F., Tarimo, E, Bodensteiner, B. L., Uyeda, J.C., & Muñoz, M. M. (2024). Opportunity begets opportunity to drive macroevolutionary dynamics of a diverse lizard radiation. Evolution Letters, In press, doi: https://doi.org/10.1093/evlett/qrae022

Eizirik, E, Murphy, W.J., Koepfli, K.-P., Johnson, W. E., Dragoo, J.W., Wayne, R.K., & O'Brien, S. J. (2010). Pattern and timing of diversification of the mammalian order Carnivora inferred from multiple nuclear gene sequences. Molecular Phylogenetic and Evolution, 56(1), 49-63. doi: https://doi.org/10.1016/j.ympev.2010.01.033

Jones, K. E., Bielby, J., Cardillo, M., Fritz, S. A., O'Dell, J., Orme, C. D. L., Safi, K., Sechrest, W., Boakes, E. H., Carbone, C., Connolly, C., Cutts, M. J., Foster, J. K., Grenyer, R., Habib, M., Plaster, C. A., Price, S. A., Rigby, E. A., Rist, J., Teacher, A., Bininda-Emonds, O. R. P., Gittleman, J. L., Mace, G. M., and Purvis, A. 2009. PanTHERIA: a species-level database of life history, ecology, and geography of extant and recently extinct mammals. Ecology, 90(9), 2648. doi: https://doi.org/10.1890/08-1494.1

O'Meara, B. C., Ané, C., Sanderson, M. J., & Wainwright, P. C. (2006). Testing for different rates of continuous trait evolution using likelihood. *Evolution, 60(5)*, 922-933. doi: https://doi.org/10.1111/j.0014-3820.2006.tb01171.x

Pennell, M. W., Eastman, J. M., Slater, G. J., Brown, J. W., Uyeda, J. C., FitzJohn, R. G., Alfaro, M. E., & Harmon, L. J. (2014). geiger v2.0: an expanded suite of methods for fitting macroevolutionary models to phylogenetic trees. Bioinformatics, 30(15), 2216-2218. doi: https://doi.org/10.1093/bioinformatics/btu181

Redding, D. W., DeWolff, C. & Mooers, A. O. (2010). Evolutionary distinctiveness, threat status and ecological oddity in primates. Conservation Biology, 24(4), 1052-1058. doi: https://doi.org/10.1111/j.1523-1739.2010.01532.x

Slater, G.J., Harmon, L. J., & Alfaro, M. E. (2012). Integrating fossils with molecular phylogenies improves inference of trait evolution. Evolution, 66(12), 3931-3944. doi: https://doi.org/10.1111/j.1558-5646.2012.01723.x

Sylvestro, D., Kostikva, A., Litsios, G., Pearman, P. B., & Salamin, N. (2015). Measurement errors should always be incorporated in phylogenetic comparative analysis. *Methods in Evolution and Evolution, 6(3)*, 340-346. doi: https://doi.org/10.1111/2041-210X.12337

Vos, R.A., & Mooers, A. O. (2006). A new dated supertree of the Primates. Chapter 5. In: Vos, R.A. (Ed.) *Inferring large phylogenies: the big tree problem*. [Ph.D. thesis]. Burnaby BC, Canada: Simon Fraser University.

Wozencraft, W. C. 2005. Order Carnivora. In Wilson, D. E. & Reeder, D. M. (Eds.) *Mammal Species of the World*. Johns Hopkins University Press.
